# Dissociable neural circuits underlie the resolution of three discrete sources of competition during task-switching

**DOI:** 10.1101/581777

**Authors:** Kelly M. Burke, Sophie Molholm, John S. Butler, Lars A. Ross, John J. Foxe

## Abstract

Humans perform sub-optimally when juggling more than one task, but are nonetheless required to multitask during many daily activities. Rapidly and effectively switching attentional focus between tasks is fundamental to navigating complex environments. Task-switching paradigms in conjunction with neuroimaging have identified brain networks underpinning flexible reallocation of cognitive resources and a core network of neural regions is repeatedly implicated (i.e., posterior parietal, inferior frontal, anterior cingulate, and middle frontal cortex). Performance costs such as reduced accuracy and slowed responses accompany the first execution of a task following a task-switch. These costs stem from three main sources of competition: 1) the need to reconfigure task-rules, 2) the immediate history of motor responding, and 3) whether inputs to be acted upon provide congruent or incongruent information regarding the appropriate motor response, relative to the recently “switched-away-from” task. Here, we asked whether both common (domain-general) and non-overlapping (dissociable) neural circuits were involved in resolving these three distinct sources of competition under high-demand task-switching conditions. Dissociable neural circuits were active in resolving each of the three sources of competition. No domain-general regions were implicated in all three. Rather, two regions were common across rule-switching and stimulus incongruence, and five regions to incongruence and response-switching. Each source of conflict elicited activation from many regions including the posterior cingulate, thalamus, and cerebellum, regions not commonly implicated in the task-switching literature. These results suggest that dissociable neural networks are principally responsible for resolving different sources of competition, but with partial interaction of some overlapping domain-general circuitry.

## Introduction

The mechanisms by which cognitive control is exerted in the human brain are of fundamental interest to neuroscientists. These neural processes are integral to our ability to flexibly change what we attend and respond to from moment-to-moment. When these processes are dysfunctional, such as in Autism Spectrum Disorder (de Vries & Geurts, 2012; Reed & McCarthy, 2012; Chmielewski & Beste, 2015) or schizophrenia (Lesh *et al*., 2011; Dickson *et al*., 2016), they can greatly impact an individual’s ability to successfully navigate daily life. One classic and highly effective way to study cognitive control is to ask experimental participants to switch from one task to another and compare their performance during this switching to when they simply repeat either of the assigned tasks alone. It is well-established that this switching incurs costs: response speeds are typically slower and task accuracy poorer following a task-switch than following a task-repeat (Jersild, 1927; Spector & Biederman, 1976; Rogers & Monsell, 1995; Wylie *et al*., 2003a; Koch & Allport, 2006; Foxe *et al*., 2014; Weaver *et al*., 2014). A considerable body of work has revealed a set of key contributing factors to these costs. 1) The first reason is the most intuitive: getting rid of the rules held in the brain for task A and loading up the rules for task B likely incurs a time cost. We will refer to this here as the *rule-switch* cost. 2) Second, research indicates that motor response history also matters. That is, if the previous task required response X but the current one requires response Y, regardless of whether the task changed or not, this *response-switch* generally incurs a cost. 3) Third, evidence shows that rules from the previous task cannot be completely inhibited, such that if stimuli or a stimulus dimension from the previous task are still present, they interfere with performing the current one. This stimulus-based interference will be referred to as a *stimulus incongruence* cost for our purposes going forward. Thus, there are three known sources of “competition” that contribute to switch costs; competition in the sense that competing task rules, competing response histories, and competing stimulus-response mappings all play their part in slowing the task-switching process and in reducing the accuracy of performance.

A key question is how each of these sources of competition is resolved in the brain: what are the underlying neural circuits, and are each resolved through activation of common or partly dissociable neural circuits? Although previous work has been done in this field, no single study has investigated the underlying circuitry of all three sources of competition: *rule-switching*, *response-switching*, and *stimulus incongruence*. Thus, a precise delineation of the subcomponents of the neural reconfiguration processes that regulate cognitive control during task-switching has yet to be conducted. There is precedence in the behavioral literature that while these three aspects of cognitive control interact with each other, they can be independently manipulated and thus represent dissociable cognitive processes (Goschke, 2000; Meiran *et al*., 2000; Wylie *et al*., 2004b).

Most prior cognitive flexibility studies have focused on examining the rule-switching aspect of task-switching, and have reliably demonstrated that regions in the frontal and parietal lobes are more active during a switch of rule than during a task repetition (Dove *et al*., 2000; Braver *et al*., 2003; Wylie *et al*., 2003b; 2004a; Buchsbaum *et al*., 2005; Liston *et al*., 2006; Wylie *et al*., 2009; Greenberg *et al*., 2010). These commonly active regions include the posterior parietal cortex (PPC; all acronyms used in this paper can be found in Supplementary Table A), inferior frontal junction (IFJ), pre-supplementary motor area (pre-SMA), middle frontal cortex, and anterior cingulate cortex (ACC). Nonetheless, not all rule-switching studies have found all of these regions to be active. This is likely due to the variety of tasks that have been used across studies, which can be addressed in part through meta-analysis to determine the regions consistently involved in cognitive flexibility. One such meta-analysis of five types of switching (shifting between rules, objects, attributes of objects, stimulus-response mappings, or locations) addressed which, if any, regions were involved in all five types of switching, and thus could be interpreted as generalized cognitive control regions (Wager *et al*., 2004). They identified seven such regions - medial prefrontal cortex, right pre-SMA, left and right anterior and posterior intraparietal sulcus, and left inferior temporal cortex – while a subsequent meta-analysis determined that the left inferior frontal junction and left posterior parietal cortex were the common regions involved in three types of rule-switching (switching between perceptual features, stimulus-response mappings, or between task rules) (Kim *et al*., 2012). Thus, across a variety of task-switching studies, the parietal lobe and frontal regions such as the inferior frontal junction and medial prefrontal cortex were consistently engaged (Vallesi *et al*., 2015).

Within a task-switching study, when the behavioral response on the current trial requires a response that is different from the previous trial, there is typically a slowing down in response time due to this response-switching (Meiran, 1996; Meiran *et al*., 2000; Mayr, 2002; Wylie *et al*., 2004b; Dreisbach *et al*., 2006). However, response-switching and rule-switching interact in an interesting way. When response-switch trials are separated by whether they were a rule-switch trial or rule-repeat trial, participants are faster when there is a rule-switch compared to a rule-repeat. On the other hand, separating out response-repeat trials in the same way demonstrates that rule-repeating is faster than rule-switching (Wylie *et al*., 2004b). Overall, it is easier when both modalities switch or when both repeat, rather than when one switches and the other repeats. In Go/No-Go paradigms, a response to a trial may even be necessary to elicit the rule-switch cost in the first place (Schuch & Koch, 2003). However, we are aware of no study that explicitly investigated the neural sources of the response-switch cost, despite a need to better understand which regions are active during these distinct sources of competition during task-switching (Wylie *et al*., 2004b).

Typical task-switching experiments require that the participant perform two different tasks on the same kind of stimulus (e.g. a colored shape). One task may require the participant to respond based on the color of the stimulus while the other task requires responding purely based on the shape of the stimulus. Responding correctly, therefore, requires attending to the relevant stimulus parameter and ignoring the other. However, when each stimulus dimension indicates a different response, this stimulus incongruence typically slows participants compared to when each stimulus dimension indicates the same response (Meiran, 1996; Mecklinger *et al*., 1999; Meiran *et al*., 2000; Mayr, 2002; Mayr *et al*., 2003; Wylie *et al*., 2004b; Dreisbach *et al*., 2006; Wylie *et al*., 2006). The stimulus incongruence cost can also be augmented when the current task is a rule-switch. (Rogers & Monsell, 1995; Goschke, 2000; Meiran, 2000; Waszak *et al*., 2005). On this kind of trial, the irrelevant stimulus dimension was also the most recently relevant one and is now indicating a different response than the current relevant dimension, possibly making the irrelevant stimulus dimension even more distracting. Studies investigating the neural sources of stimulus incongruence costs generally implicate the ACC (Mayr *et al*., 2003; Swainson *et al*., 2003; Fassbender *et al*., 2004; Woodward *et al*., 2006; Kim *et al*., 2010). One theory about the ACC’s role in stimulus incongruence costs posits that the ACC is involved in resolving conflict of information-processing, which subsequently activates the prefrontal cortex – a region commonly implicated in task-switching studies - to reduce the conflict susceptibility (Botvinick *et al*., 2001). More specifically, evidence indicates that the ACC may register a change in conflict that needs to be resolved (Mayr *et al*., 2003). In addition to ACC, two specific regions likely to be engaged by manipulations of stimulus incongruence are PPC, where activity has been shown to increase with the salience of the irrelevant stimulus (Liston *et al*., 2006), and the inferior frontal junction which appears to be involved in cognitive control, particularly in studies using strongly incongruent stimuli that are difficult to ignore (Derrfuss *et al*., 2004; Derrfuss *et al*., 2005).

Given the dearth of studies on the neural sources of *response-switching* and *stimulus incongruence* specifically, we can take advantage of the extensive task-switching literature and conflict monitoring literature to predict a subset of frontal and parietal regions to be involved in each of the three sources of competition involved in task-switching. Here, we utilized a multi-session approach to acquire at least four times as much data per participant as is typical to gain intra-individual reliability and sensitivity with a repeated measures analysis. We hypothesize that each source of competition will reveal engagement of unique, dissociable neural circuits that are specific to that source of competition, as well as domain-general regions active in resolving more than one source of competition. For example, rule-switching may necessitate activation of the pre-SMA because of the changing association of abstract rules with appropriate motor responses (Bunge *et al*., 2003; Boettiger & D’Esposito, 2005), while response-switching may engage inferior frontal cortex, a region involved in regulating response inhibition (Rubia *et al*., 2001; Bell *et al*., 2014a; Bell *et al*., 2014b; Scalzo *et al*., 2016). Moreover, stimulus incongruence may increase activity in the ACC, a region involved in conflict detection (Mayr *et al*., 2003; Swainson *et al*., 2003; Fassbender *et al*., 2004; Woodward *et al*., 2006; Kim *et al*., 2010), and IFJ which is active when salient, behaviorally-relevant stimuli are present (Corbetta & Shulman, 2002). At the same time, we predict that all three sources of competition will involve activation of the generalized cognitive control regions PPC and prefrontal cortex (e.g. IFJ).

## Methods

### General Approach

We employed a multi-session approach whereby participants performed a task-switching paradigm over four separate functional magnetic resonance imaging (fMRI) scanning sessions (i.e., across four separate testing days), with the principle aim of this approach being to greatly enhance power at the individual participant level. All neuroimaging analyses used a repeated measures approach to account for the multi-session data. It should be noted that all participants also completed four additional sessions while high-density electroencephalography (EEG) data were recorded. These data are not reported here and will be reported separately. The Institutional Review Board of Albert Einstein College of Medicine approved all materials and procedures, and all ethical guidelines were in accordance with the tenets of the Declaration of Helsinki.

### Participants

Eleven adults with normal or corrected-to-normal vision participated. Individuals were excluded from the study if they had metal in their bodies, a history of brain injury, or a psychiatric diagnosis (see Table 1). One participant was not able to attend the fourth fMRI session, and one session from another participant needed to be excluded due to movement artifact; therefore, there were 42 experimental sessions (days) from eleven participants included in all analyses. Participants provided informed, written consent and were modestly compensated for their time.

**Table 1.**
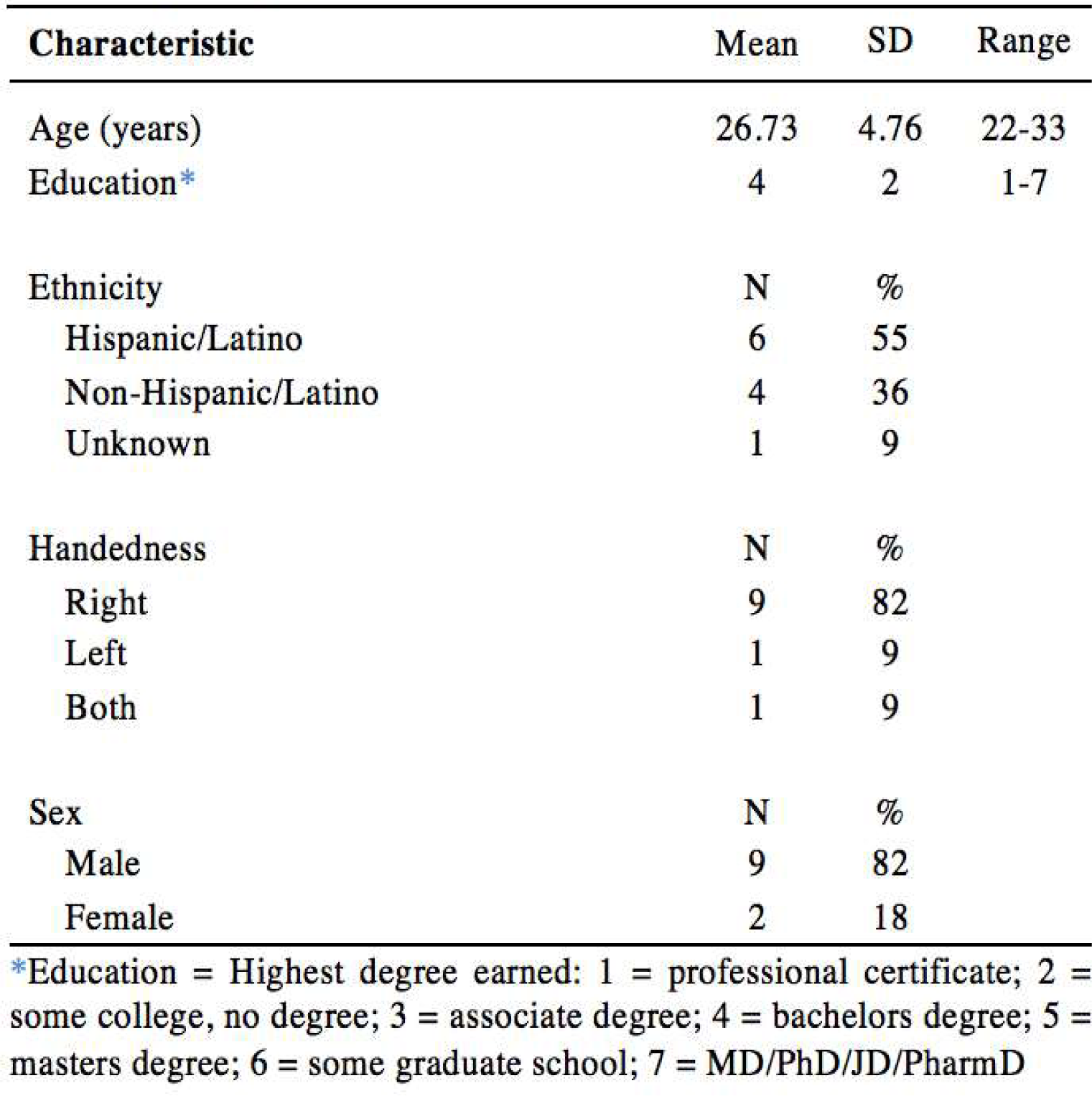
Demographics of Sample (N=11)

### Stimuli and Task

A cued, speeded, task-switching paradigm was presented using Presentation software (NeuroBehavioral Systems) which was based on a design to isolate rule-switching preparatory processes from stimulus decoding and motor response processes (Spector & Biederman, 1976; Wylie *et al*., 2004b; 2006). Throughout the entirety of the experiment, the monitor displayed a gray background of [128 128 128]. Each two-second long trial commenced with a colored rectangle subtending 4.25° vertically and 6.38° horizontally of visual angle in the center of the screen for 200 ms, which was the task cue (i.e. yellow for the letter task, blue for the number task; this color-task association was randomly assigned for each participant and remained constant for all four sessions). This was followed by small central crosshairs for 500 ms and then the imperative stimuli appeared for 200 ms, which were a letter and number randomly assigned to either side of the crosshairs each subtending 1.18° vertically (see Figure 1). A screen with crosshairs was displayed for the remaining 1100 ms while the participant responded or withheld a response with a button press on a response pad. When the cue indicated the letter task the participants were instructed to “respond as quickly and accurately as possible to the letter task by responding with a button press for a vowel and withholding for a consonant. For the number task, respond as quickly and accurately as possible by responding with a button press for even numbers and withholding for odd numbers.” Letters were randomly selected from the set [A E G I K M R U] and the numbers were randomly selected from the set [2 3 4 5 6 7 8 9]. Each trial contained both a letter and number so the task cue was integral to responding appropriately to the stimulus. The side of the crosshairs to which the letter and number were displayed was random so that participants could not anticipate which side of the screen to attend to. Given the random selection of stimuli, rule-switches were 50% probable; incongruent stimuli and congruent stimuli were equally probable; and response-switches were equally likely as response-repeats. Four trials were presented in a row followed by six seconds of rest to allow for BOLD signal decay. This trial sequence was repeated 16 times within a block, after which participants could rest before starting the next block, and 16 blocks were run per session. Four blocks were Pure blocks where only one task was cued. Overall this means that there were 2304 trials included per participant for all analyses. This was important to ensure the participant’s ability to perform each task on its own. If there were any changes in behavior throughout the session, we could determine if this was due to the switching of tasks or a general fatigue. Whether a participant started the session with the letter task or number task was counterbalanced between sessions and between participants. At the beginning of the participant’s first session, they practiced ten trials of the letter task and ten trials of the number task to ensure they understood the experiment and its timing.

**Figure 1.**
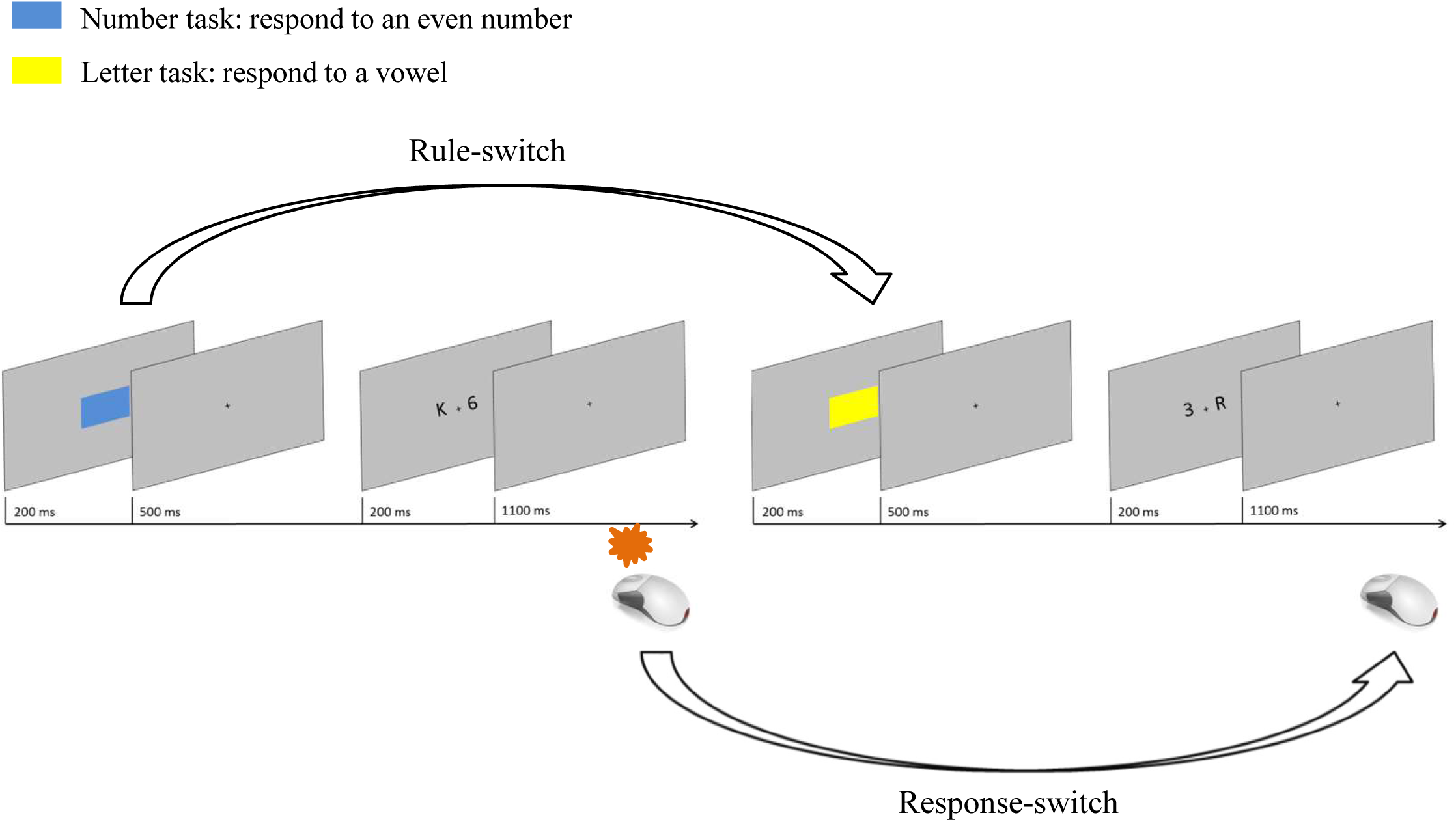
Example stimuli for two trials: a color cue was presented to indicate the task, 700 ms later the stimuli appeared and the participant responded within 1300 ms. The next color cue appeared, which could switch or repeat rules, thus creating a rule-switch or rule-repeat trial. Trials could have stimuli which were congruent with each other (i.e. indicating the same response for the letter task and number task) or incongruent with each other (i.e. the letter and number indicating different responses). In addition, each trial could be categorized by whether the motor response changed from the previous trial (e.g. a No-Go trial followed by a Go trial) or whether it repeated (e.g. a No-Go trial followed by a No-Go trial).

### Condition Definitions

The first trial from each four-trial series was excluded as it represented neither a rule-switch nor rule-repeat. A rule-switch trial was where the cued task was different from the previous (e.g. the letter task preceded by the number task or vice versa), and a rule-repeat trial was where the same task was cued as in the previous trial. Response-switch trials required a different response (either withholding a response or making a response) from the previous trial whereas response-repeat trials called for the same response as in the preceding trial. Response time analysis was necessarily constrained to those response-switch or response–repeat trials involving a motor response, but the fMRI analysis could take advantage of both successful withhold trials and successful response trials. Stimulus incongruent trials were those in which one stimulus indicated to respond and the other indicated to withhold a response (e.g. a vowel and odd number, or a consonant and even number). Stimulus congruent trials had stimuli that both indicated the same response (e.g. a vowel and even number, or a consonant and odd number).

### Behavioral Analysis

Response times (RTs) were calculated for each session of a given participant according to the three sources of “conflict” (i.e. rule-switching, stimulus incongruence, and response-switching). Accuracy was determined as the percentage of correct responses (correct hits + correct rejections) to the total number of trials (correct hits + correct rejections + omission errors + commission errors). Behavioral costs for each participant were calculated according to each of the three conflicts (e.g. rule-switch cost = average rule-switch RT – average rule-repeat RT). If two button press responses occurred within one trial, only the first response was included in analysis; the second one was interpreted as an extraneous response. Minimum and maximum response times were determined by excluding any responses that occurred within 200 ms of the presentation of the stimulus (these responses are seen as either late responses to the previous trial or extraneous mouse clicks), and excluding responses that occurred more than 2.5 standard deviations after the mean response time for that session (Manzi *et al*., 2011). An omnibus 4-way ANOVA on the three conditions of interest and session (first, second, third, fourth) was conducted on the response times to determine whether the factors of interest interacted, and to verify whether repeated sessions had any effect on the data as well.

### fMRI Acquisition

Participants lay comfortably supine in a 3.0 T Philips TX Achieva MRI scanner (Royal Philips, Amsterdam, Netherlands), with foam earplugs and heavy-duty headphones over them, and with goggles laying gently over their eyes to display the computer screen (using Resonance Technology VisuaStimDigital with funclab software). The scanner has a thirty-two channel head coil and acquired T2, EPI, FLAIR, DTI, and T1-weighted structural MPRAGE (Magnetization Prepared Rapid Gradient Echo) scans. Whole-brain functional scans were acquired through twelve blocks of 114 volumes. Each volume consisted of forty-eight axial slices in an ascending interleaved order at 3 x 3 x 3 mm³ using a T2-weighted echo-planar sequence (TR/TE: 2000/20 ms, flip angle = 90°, 240 mm FOV). The short TE was used because we were specifically interested in ensuring good signal capture from anterior cognitive control regions where there are usually field inhomogeneities (Weiskopf *et al*., 2006; Weiskopf *et al*., 2007). We fully expected that anterior regions would be involved in task-switching and thus wanted to capture these regions well, and although we were aware that the short TE sacrifices BOLD signal slightly, the long and repeated scan sessions (3.5-4 hours per participant) was expected to compensate for this. The inter-stimulus interval was 2000 ms, as well as the TR, so oversampling was not employed in the current study. FMRI sessions were a total of 1 hour and 20 minutes, and the task was presented for 60 minutes of that time.

### fMRI Preprocessing

FMRI processing was carried out using BrainVoyager QX v2.4 (Brain Innovation BV, The Netherlands) (Goebel *et al*., 2006). The T1-weighted anatomical scans for each participant were normalized into Talairach space using the AC-PC landmark and fitting 6 parameters (the superior, inferior, anterior, posterior, left, and right-most parts of the brain). Processing for all functional runs included: removal of the first two volumes to allow for equilibrium effects, slice scan time correction, 3D motion correction using a trilinear/sinc interpolation and 6 vectors (translation in the X, Y, and Z dimensions, 3 Euler angles of rotation), temporal filtering (high-pass GLM-Fourier basis with 2 sines/cosine), normalization to Talairach space using the AC-PC landmark and fitting 6 parameters), and spatial smoothing using an 8 mm isotropic full-width half-maximum Gaussian filter kernel. Each functional run was precisely aligned and co-registered to the T1 anatomical scan. Any runs with excessive movement (greater than 3 mm of translation or 3° of rotation) were excluded. This excluded one scan session of one participant, which had too much movement. In addition, only 3.9% of all remaining runs from all participants were excluded due to excessive motion.

### fMRI Activation Analyses

An event-related design was used to examine the three sources of conflict present in this experiment: rule-switching (rule-switch > rule-repeat), response-switching (response-switch > response-repeat), and stimulus incongruency (incongruent stimuli > congruent stimuli). Each condition of interest (rule-switch, rule-repeat, response-switch, response-repeat, incongruent stimuli, congruent stimuli) was modeled with a boxcar function that was convolved with a standard two-gamma hemodynamic response function (with an event length assumed to be 2000 ms, 5 seconds until response peak, 15 seconds until the undershoot peak, and a response undershoot ratio of 6). Volume time-courses for each run of each participant were entered into a random-effects general linear model (RFX-GLM) in BrainVoyager, which was used in an ANCOVA Random Effects Analysis. The ANCOVA was set up with two within-subjects factors: 2 conditions x 4 sessions. This approach followed done separately and independently for each of the three sources of conflict (rule-switching vs rule-repeating, incongruency vs congruency, and response-switching vs response-repeating). The ANCOVA tested for the Condition factor, resulting in a map where each voxel had an F-stat and its associated p-value. fMRI analysis inherently has multiple statistical comparisons and so thresholding of some kind is required to statistically compensate for increased false positives. For each of the three contrasts of interest (rule-switching, incongruency, and response-switching) we applied p-value- and cluster-thresholding. The map’s p-value threshold was decreased to below 0.01 and then the Cluster-Level Statistical Threshold Estimator in BrainVoyager was used. In order to correct for multiple statistical comparisons, this estimator is based on a Monte Carlo simulation of random image generation, with added spatial correlations between neighboring voxels, voxel intensity thresholding, and cluster identification. After 1,000 simulations, a minimum cluster size was determined for that statistical map that yields a false positive detection rate of 5% or less (Forman *et al*., 1995). The cluster threshold estimator determined that cluster minimum should be 30 voxels for the rule-switching data, 18 voxels for the incongruency data, and 24 voxels for the response-switching data. These thresholds were used for each of the maps. The average beta-weight for the voxels within each contiguous region for each condition was calculated and plotted to show the absolute difference in activation or deactivation. The centers of mass in Talairach coordinates were used to determine the brain region that best represented the region of activation, according to the Talairach Daemon (http://www.talairach.org/).

### Overlapping Regions

To determine which regions were involved in more than one source of conflict, we applied conjunction analysis in BrainVoyager to map voxels that were significant in more than one analysis and whether any voxels were significantly involved in all three analyses.

### Effects of Time

Participants coming in for multiple sessions allows an investigation of whether there are changes in behavior or network activity across time. To determine if there were any effects of session on the response time data, we used RFX-GLM in the ANCOVA Random Effects Analysis with two within-subjects factors but instead tested for the Session factor. To anticipate the results, there were no significant effects of session in the data.

## Results

### Behavioral Performance

The three-way ANOVA demonstrated an interaction between the three sources of conflict of interest (F(1,10) = 9.866, p = 0.010) and a 2-way interaction between rule-switch and response-switch (F(1,10) = 9.342, p = 0.012). Post-hoc t-tests revealed that response-repeating was faster with a rule-repeat than a rule-switch (*t*(21) = 2.83, p = 0.010) and response-switching was faster with a rule-switch than with a rule-repeat but this was not significant (*t*(21) = −1.55, p = 0.136). Interestingly, the expected 2-way interaction between rule-switch and stimulus incongruence was not significant (F(1,10) = 3.818, p = 0.079). In addition, although the average response times indicated trends toward rule-switch costs, stimulus incongruence costs, and response-switch costs, these were not strong enough to emerge as main effects (see Figure 2A). Table 3 lists the average response times and standard deviations for each participant. In a study design such as this, one concern is that response times may slow to the benefit of accuracy. But in fact, the data demonstrated a correlation where slower participants were less accurate (r = −0.62, p = 0.0412; see Figure 2C). However, with a cohort of 11 participants, any correlational observations should be interpreted with caution.

**Figure 2.**
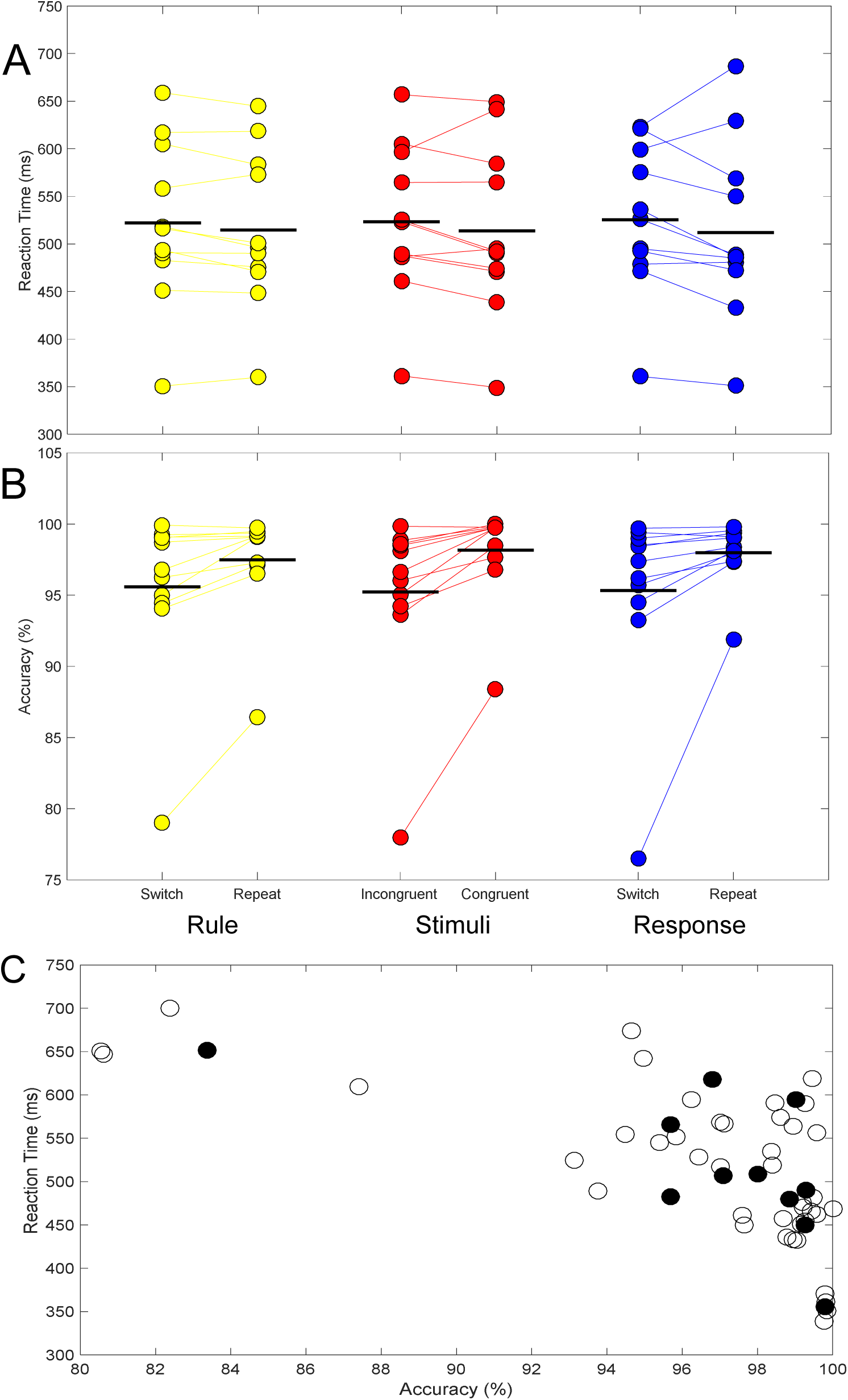
A) The average reaction times were slower for each of the three sources of conflict; B) the average accuracy was also slower; and C) reaction time and accuracy were negatively correlated based on both the averages of the participants (filled circles), or averages from each session (open circles).

**Table 2.**
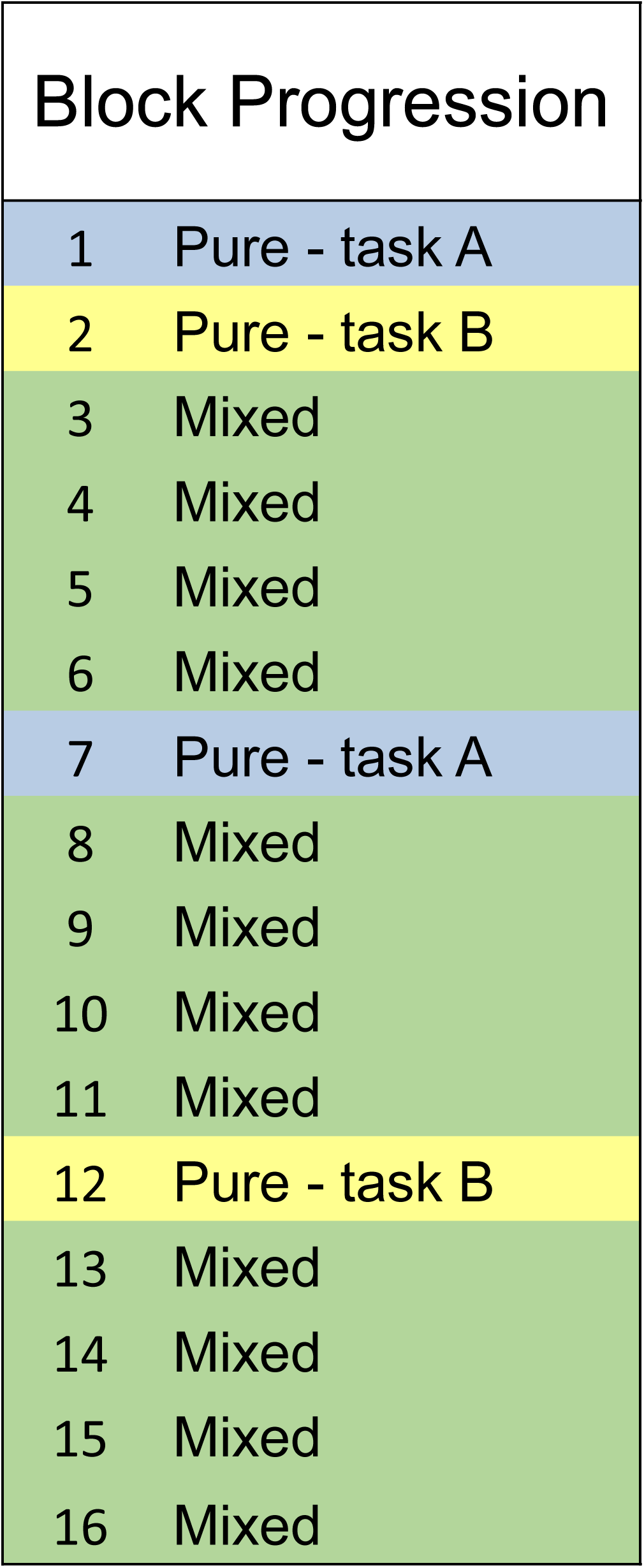
Sequence of blocks throughout one scan session. Participants started with a pure block of one task, either the letter or number task, which was counterbalanced between participants and scans. This was followed by a pure block of the other task to counterbalance experience. Systematically throughout the scan, they had two more pure blocks. In the event participants’ performance decayed over time, it would be possible to determine if it was an overall effect possible due to fatigue or a switch-specific effect.

**Table 3.**
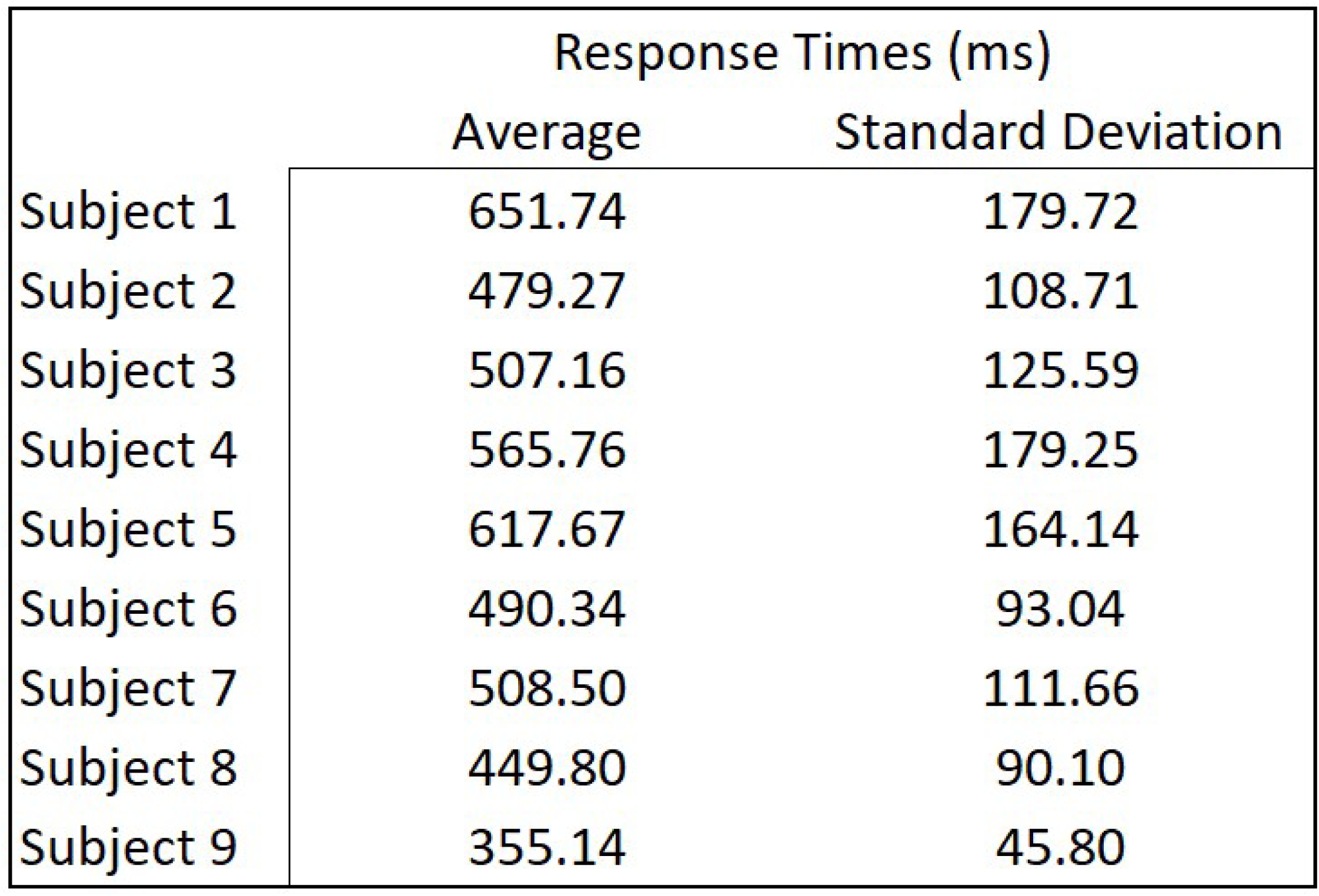
Overall average response times for each participant and their standard deviations.

### fMRI Data

The three contrasts revealed a variety of distinct regions and overlapping regions of involved in resolving the three sources of conflict (see Table 4). The contrast examining rule-switching effects revealed five regions that were significantly differentially active for rule-switch versus rule-repeat trials, that included left inferior frontal junction, left precuneus, left thalamus, and a locus in posterior cingulate that was more active for rule-switching than rule-repeating and a locus in posterior cingulate that had less deactivation to rule-switching (see Figure 3). The contrast examining incongruency effects demonstrated activations in the cingulate, superior frontal gyrus, medial frontal gyrus, middle frontal gyrus, precuneus, cerebellum, thalamus, and insula (see Figure 4). Although our analysis does not allow for investigating statistical interactions, we can still determine that two of these regions – precuneus and thalamus – were involved in both rule-switching and incongruency. Investigating response-switching effects revealed ten regions significantly involved, which were predominantly cerebellar and subcortical, along with the middle frontal gyrus, postcentral gyrus and medial frontal gyrus (see Figure 5). Interestingly, some of these regions were common to both incongruency and response-switching, with just the caudate, thalamus, postcentral gyrus, and anterior cingulate cortex unique to the response-switching contrast and did not come out in the other analyses (see Figure 6). Also, the medial frontal gyrus, right postcentral gyrus and anterior cingulate showed stronger deactivation patterns for response-switching than response-repeating. The medial frontal gyrus which was involved in resolving incongruency as well, was similarly more deactivated for the trials with incongruent stimuli compared to those with congruent stimuli.

**Table 4.**
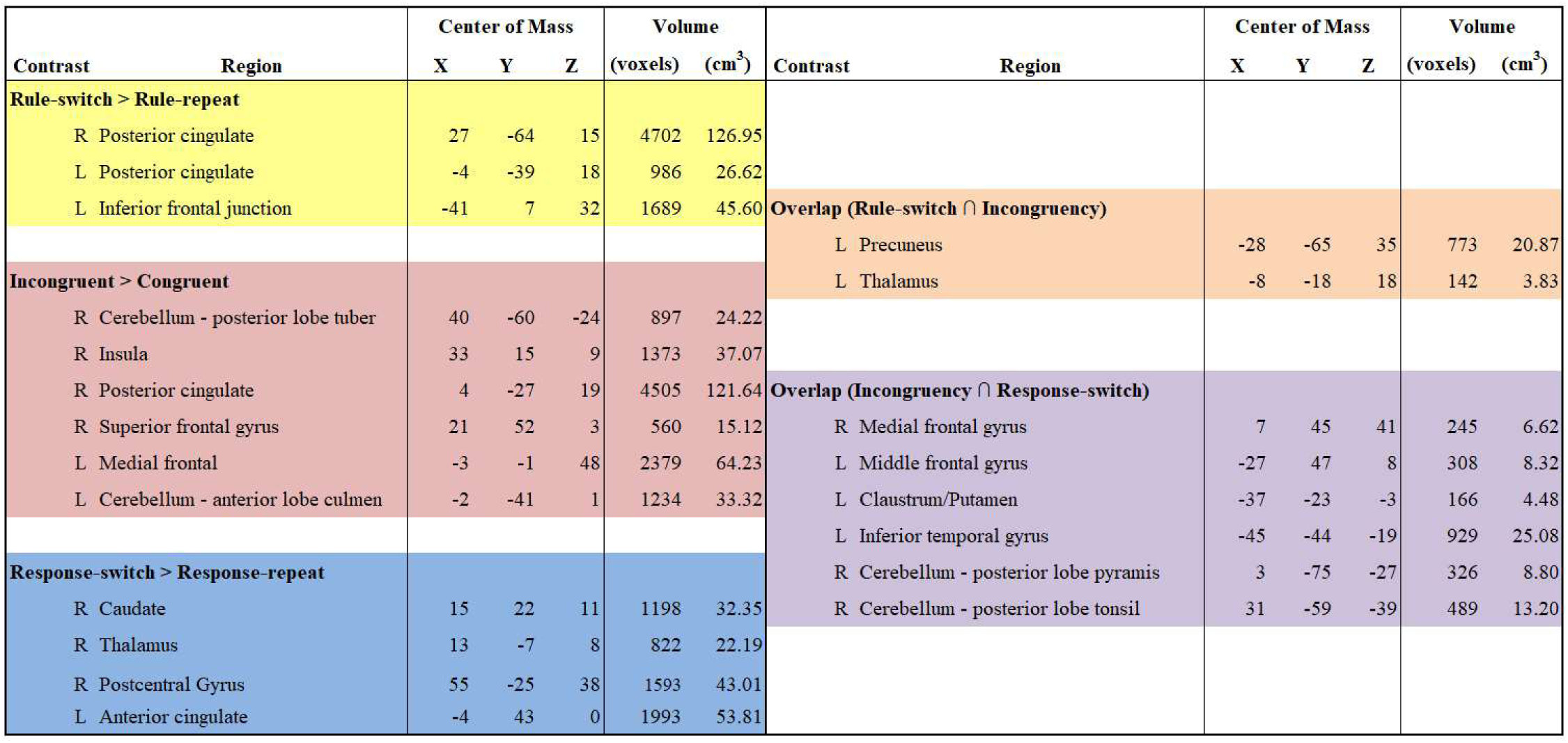
Regions that came out significant in our repeated measures ANOVA of the imaging data for task-switching, incongruency, response-switching. The left-hand side of the table shows which regions came out only for each of the three analyses. On the right-hand side of the table are the clusters of voxels that were significant in more than one analysis. Centers of mass are in Talairach coordinates. The overall average beta-weights for each contrast are shown next to each ROI.

**Figure 3.**
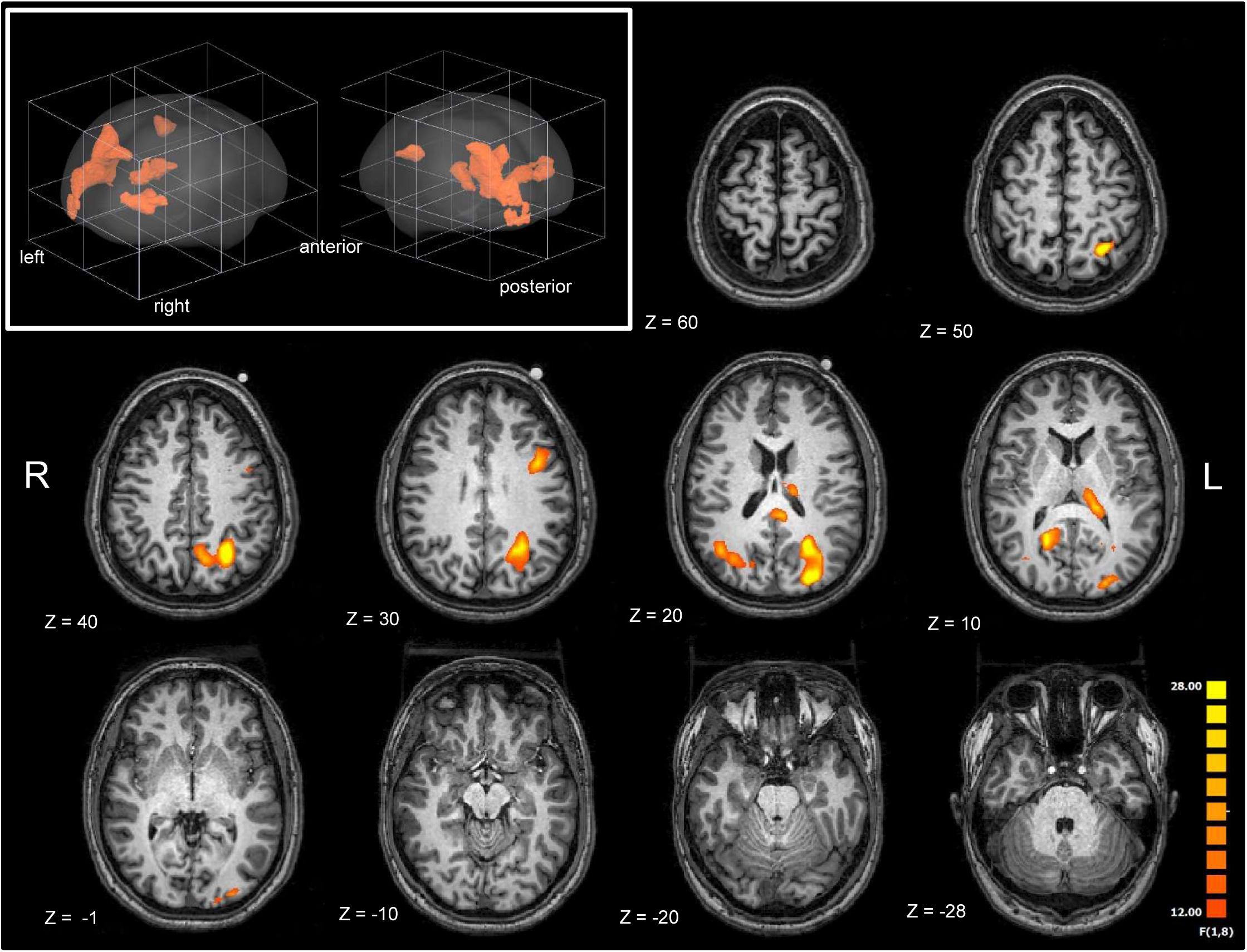
Map of regions significantly involved in rule-switching. The upper left panel shows all the significant regions in orange in a 3D rendering. In the horizontal slices, the color range is according to the F-stat for each voxel, ranging from 12 to 28. Images are in radiological format (right is left), and z coordinates are in Talairach space.

**Figure 4.**
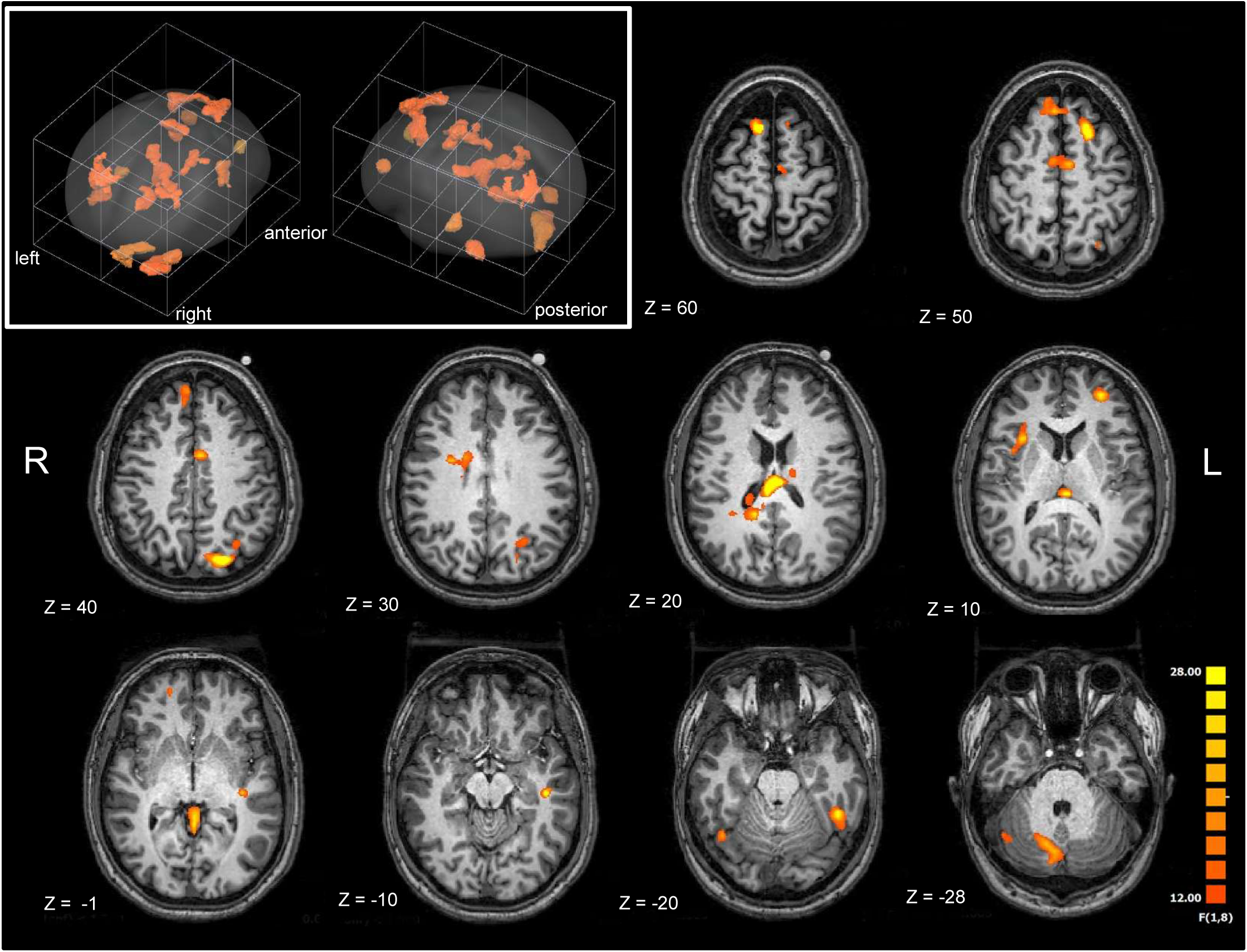
Map of regions significantly involved in incongruency. The upper left panel shows all the significant regions in orange in a 3D rendering. In the horizontal slices, the color range is according to the F-stat for each voxel, ranging from 12 to 28. Images are in radiological format (right is left), and z coordinates are in Talairach space.

**Figure 5.**
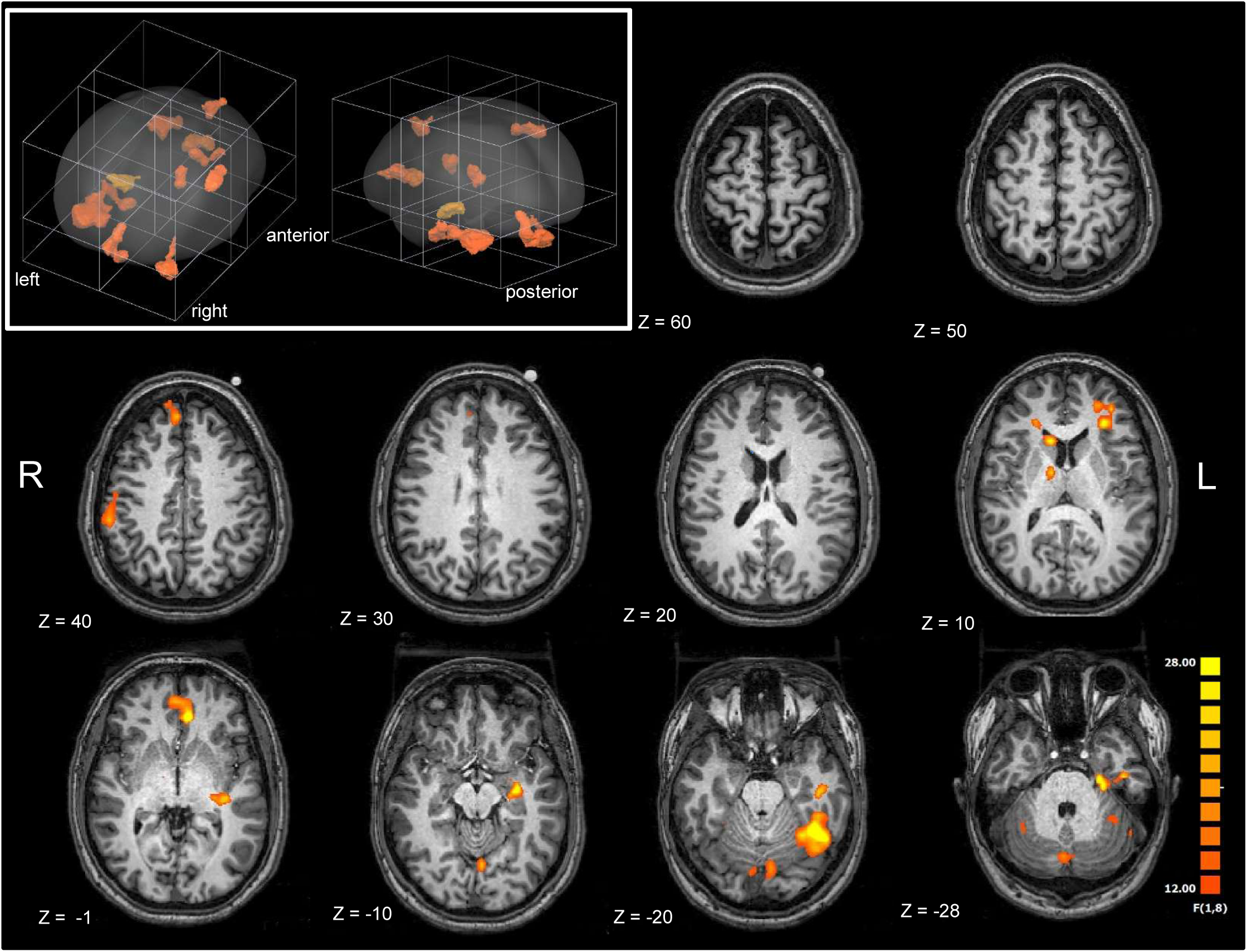
Map of regions significantly involved in response-switching. The upper left panel shows all the significant regions in orange in a 3D rendering. In the horizontal slices, the color range is according to the F-stat for each voxel, ranging from 12 to 28. Images are in radiological format (right is left), and z coordinates are in Talairach space.

**Figure 6.**
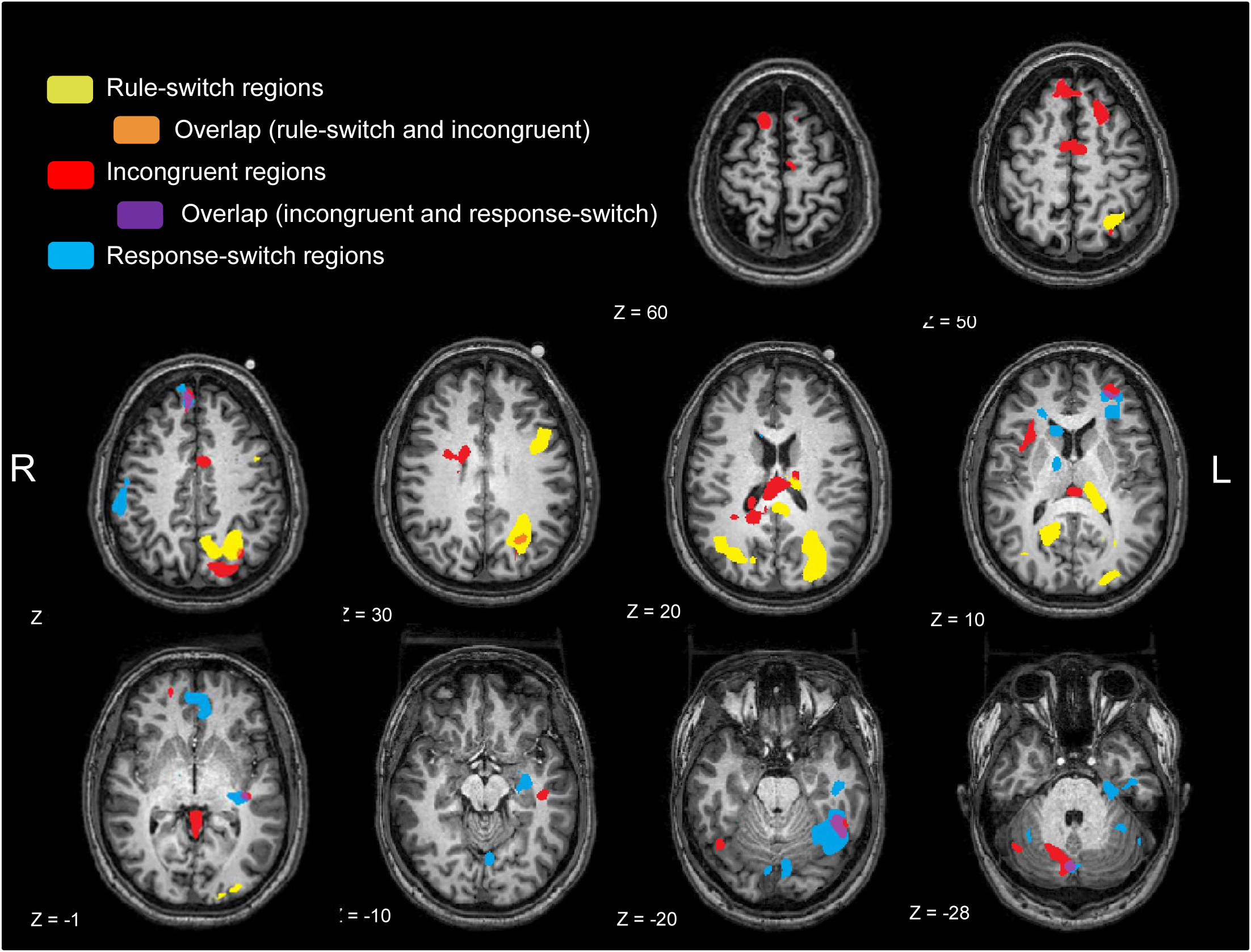
Map of all areas involved in all three sources of conflict, including where they overlap. For simplicity, solid colors are used to represent each condition or overlap of conditions. Images are in radiological format (right is left), and z coordinates are in Talairach space.

## Discussion

We set out here to map the cortical circuitry underlying three known sources of competition that contribute to response slowing during task-switching, examining the thesis that resolving these sources of conflict likely involves both domain-general cognitive control regions as well as dissociable sub-circuitry. We describe three distinct networks that were evident when the discrete sources of conflict were expressly isolated in our analyses (i.e., for rule-switching, response-switching and stimulus incongruence). Somewhat surprisingly, and contrary to our original hypothesis, there were no overlapping regions active during all three comparisons, so the current results do not support the existence of one or more domain-general cognitive control regions that subserve all three of these functions. Instead, we found that there were both frontal and subcortical regions involved in resolving conflict across a given pair of conditions, so some degree of domain generality was also observed (see Figure 7 for a schematic summary overview). The main outcome of the current study though, was the observation that largely distinct networks of regions were required to resolve each source of conflict inherent in this task-switching paradigm. Rule-switching on its own recruited the typically observed frontal and parietal regions reported in the task-switching literature (Wager *et al*., 2004; Kim *et al*., 2012) along with the thalamus and posterior cingulate cortex. Stimulus incongruence involved the cingulate as predicted but also the cerebellum and frontal regions with a medial frontal gyrus deactivation. Response-switching involved sub-cortical, anterior cingulate, and cerebellar involvement in addition to a frontal deactivation. Although all three of these analyses revealed networks of regions significantly involved, it was surprising that the behavioral analysis did not reveal main effects when each of the three sources of conflict were isolated individually. Based purely on average reaction times, our results are within the range typically found in similar task-switching studies, albeit towards the lower end. In the current cohort, response-switching elicited the largest change in reaction times, and interacted significantly with the other two sources of competition. What is striking is that despite the lack of significant rule-switch or incongruence costs, the neuroimaging data did demonstrate robust changes in neural activity. In what follows, we describe each of these extended circuits in more detail and discuss the possible roles of the various functional hubs in resolving each distinct source of competition.

**Figure 7.**
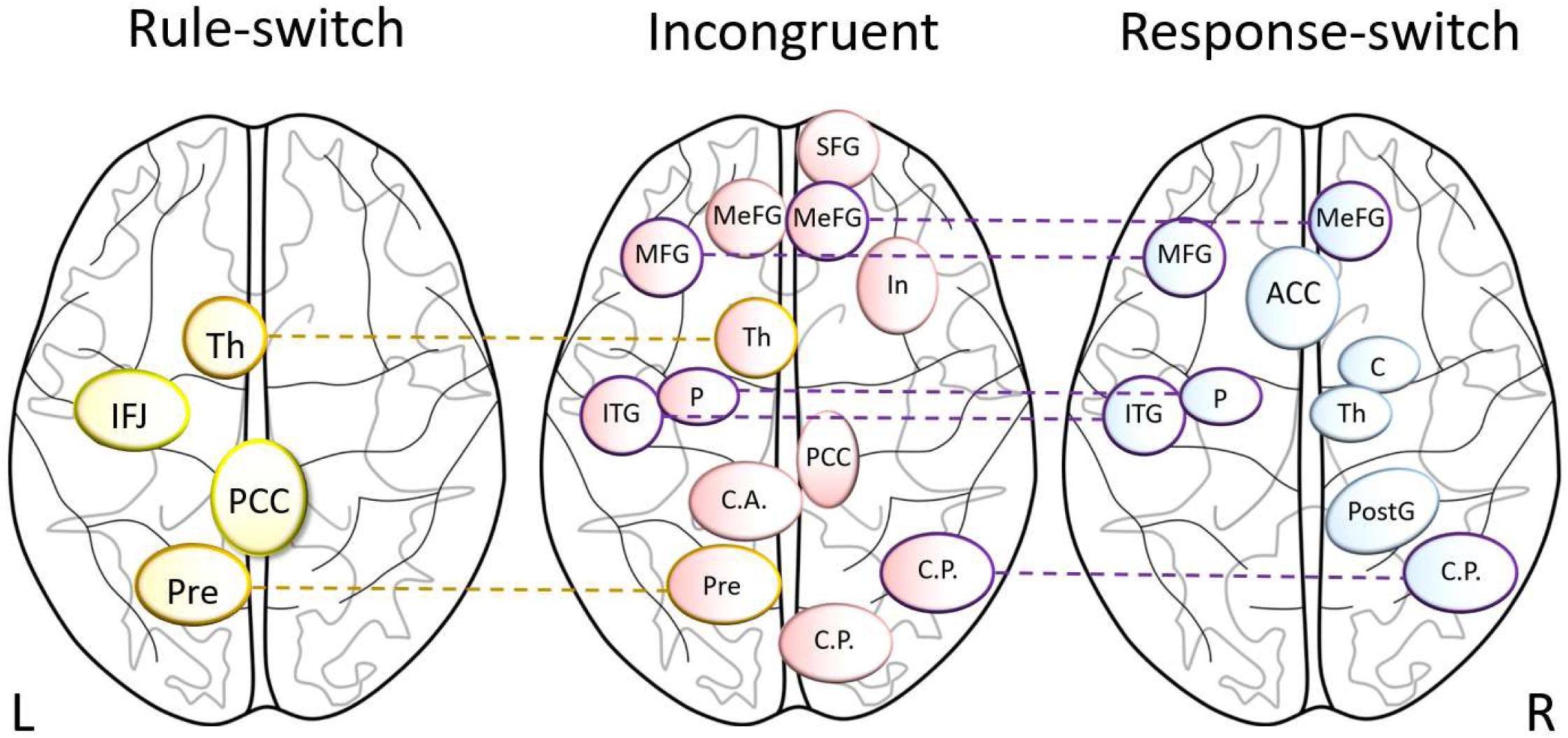
Schematic representation of the regions involved in the three sources of conflict investigated in this study. While some regions were involved in more than one source of conflict (dotted lines), there were no domain-general regions involved in all three. Legend: ACC = anterior cingulate cortex, C = caudate, C.A. = cerebellum anterior lobe, C.P. = cerebellum posterior lobe, In = insula, IFJ = inferior frontal junction, ITG = inferior temporal gyrus, MeFG = medial frontal gyrus, MFG = middle frontal gyrus, P = putamen, PCC = posterior cingulate cortex, PostG = postcentral gyrus, Pre = precuneus, SFG = superior frontal gyrus, Th = thalamus.

### Rule-switching

Rule-switching (changing the rule from letter-based to number-based or vice versa) activated the precuneus, a region consistently reported in task-switching studies. Activity in the precuneus has previously been time-locked to attention shifting in a card-sorting task (Nagahama *et al*., 1999), to attention shifting between different targets in space (Selemon & Goldman-Rakic, 1988), and has been associated with a variety of tasks compared to rest (Utevsky *et al*., 2014), so its involvement here when participants were required to reconfigure the task rule they should follow is not surprising. Similarly, the posterior parietal cortex is also consistently implicated in task-switching (see meta-amylases by (Wager *et al*., 2004; Liston *et al*., 2006; Kim *et al*., 2012)) and the current study’s finding of left posterior parietal cortex activation further confirms this region’s role in cognitive control. The clear laterality in our findings in the parietal lobe and frontal lobe are in agreement with left-laterality sometimes reported in the literature (Dove *et al*., 2000; Philipp *et al*., 2013; Muhle-Karbe *et al*., 2014). However, there is no consensus as of yet regarding which regions are bilaterally or unilaterally involved in task-switching and we will therefore discuss our findings in more general terms. Rule-switching also activated a region on the border of both the precentral gyrus and inferior frontal cortex, which is commonly referred to as the inferior frontal junction (IFJ) (Derrfuss *et al*., 2004). This region is often found to be involved in task-switching (Dove *et al*., 2000; Braver *et al*., 2003; Swainson *et al*., 2003; Brass & von Cramon, 2004). Meta-analysis has revealed the IFJ may be specifically involved in the updating of the task representation (Derrfuss *et al*., 2005). The thalamus has been shown to be involved in task-switching as well. Nuclei in the thalamus were crucial for modulating synchrony between cortical areas during an attention task in macaques (Saalmann *et al*., 2012), further indicating that the thalamus regulates information flow throughout inter-regional cortico-cortical networks (Bell & Shine, 2016).

Rule-switching was also associated with differential activation within the posterior cingulate cortex (PCC), which we had not specifically predicted. The posterior cingulate is believed to serve as a hub within the so-called default mode network, and it is thought to show generally high levels of background activity during so-called “resting states”, activity that is then significantly decreased or suppressed during task performance (Gusnard *et al*., 2001; Raichle *et al*., 2001; Hayden *et al*., 2010; Spreng *et al*., 2010; Leech *et al*., 2011; Korgaonkar *et al*., 2014). Its functional role is not yet well understood, but based on its structural and functional connections with the fronto-parietal network, it may well play a role in transitioning between network states (Hagmann *et al*., 2008; Agam *et al*., 2013) and in dynamically controlling attention from the internal to the external environment. Here, two distinct loci of activity in the posterior cingulate were significantly involved in rule-switching. One locus, which spanned 27 mm³ and was located medially, but in the left hemisphere, was not significantly activated for rule-repeating compared to baseline, but was for rule-switching. The other locus, with a center of mass in the right hemisphere that spanned 127 mm³, was significantly engaged for both conditions relative to baseline, but more so for rule-switches than repeats. These regional activations suggest a more active role for PCC in resolving rule-switching. That is, we do not observe relative decreases in activation during higher levels of task engagement. The current study is not the first to demonstrate greater PCC activity during task-switching relative to task-repeating (Braem *et al*., 2013) (Dreher *et al*., 2002; Braem *et al*., 2013). Interestingly, in the study by Braem and colleagues, PCC activity during switching was impacted by the valence of affective visual stimuli presented between trials (i.e. positive vs negative), suggesting a possible role for the PCC in the affective modulation of cognitive control processes. Indeed, a number of other studies have reported PCC involvement in evaluating the subjectivity or the salience of stimuli in general. For example, in a go/no-go task with emotionally salient faces, PCC was active during successful task performance when the participant was under conflicting emotional conditions. Participants were put under an emotional state of threat (e.g. the possibility of hearing aversive sounds) or excitement (e.g. the possibility of winning money) and on each trial an emotional cue, a happy, sad, or neutral face was presented. On trials where the emotion of the presented cue conflicted with the emotion of the overall state (e.g. threat condition and a happy face), the PCC was more active than when the emotional states did not conflict. (Cohen *et al*., 2016). Studies have also correlated activity of the PCC with the subjective value of a chosen option in studies on decision-making in both monkeys and humans (Kable & Glimcher, 2007; Luhmann *et al*., 2008; Levy *et al*., 2011). The generalizability of these affect- or salience- related findings to the current study is, of course, somewhat limited since there was no affective or salience manipulation of the stimulus materials.

A recent study provides another interesting wrinkle on the potential role of the PCC in rule switches. Manelis and colleagues presented participants with a series of objects with the task being to determine if a given object was “old” (seen before within a given block) or “new” (never seen before) (Manelis *et al*., 2017). In a variant of this task, they then presented occasional reset signals such that previously “old” objects should now be considered as “new” for the purposes of that task block – which they termed “pseudonew”. The PCC was the sole area to show complete sensitivity to this task-resetting manipulation, with its activity reflecting an item’s novelty (or not) according to the task instructions, regardless of whether the item had been seen during preceding blocks. In other words, the posterior cingulate was sensitive to the rule-switch, much as we find here.

Overall, the current findings and much of the extant literature would suggest that the posterior cingulate plays a more prominent role in task-switching than might have been previously appreciated, likely as a region that detects conflicts of many types and reallocates resources of other networks to optimize further processing. The PCC, with strong connections to parahippocampal areas, could also play a role in retrieving relevant information from long-term memory (Kim *et al*., 2010; Kim, 2011). We can only speculate at this point but further studies into changes of rules under different contexts will likely better delineate its specific role, and we will return to the potential role of the posterior cingulate below when we discuss its involvement in the resolution of stimulus-stimulus incongruence.

In sum, the regions involved in the current rule-switching findings suggest a more detailed and elaborate set of perceptual-cognitive processes during successful rule-switching than has been forwarded previously. One possible way of thinking about how these regions could work together to resolve rule-switching (although we would stress that this is but one possible sequence), could be: increased transition from internally-guided thoughts to the external environment (posterior cingulate), increased information flow between networks (thalamus), update of the task (IFJ), and increased attention to the visuospatial input (precuneus).

### Response-switching

The activations to response-switching were also revealing. A response-switch occurred when the correct response to the current trial was different from the previous trial (e.g. a Go when the previous trial was a No-Go), regardless of the task. Response-switching elicited differential activity in the left middle frontal and left inferior temporal gyrus. In addition, sub-cortical (specifically the caudate and thalamus) and cerebellar activations were also observed. A somewhat surprising finding was that the medial frontal cortex was more active for response-repeating compared to response-switching. However, in both of these conditions, activity levels were actually less than during rest (i.e. the beta-weights were negative for both conditions). Previous studies suggest that the medial frontal cortex is integral to the medial frontal-subcortical circuit which has been implicated as a mediator of motivation, engagement, and the maintenance of task-relevant processes (Stuss *et al*., 2005). As such, it is perhaps surprising that this region was not significantly active above baseline for response-switches.

Middle frontal cortex was also significantly involved in response-switching. It is noteworthy that activity levels in middle frontal cortex during response-repeating were similar to baseline, whereas they were significantly increased above baseline for response-switching. This observation accords well with meta-analysis findings, where bilateral middle frontal cortex was found to be involved in response suppression during Go/NoGo tasks, and during Wisconsin Card Sorting Tasks (WCST), but not specifically in task-switching studies (Buchsbaum *et al*., 2005). Here, we find no evidence for its involvement in rule switching, but clear involvement during response switching.

The posterior lobe of the cerebellum was also implicated in the response-switching analysis (and during analysis of stimulus-stimulus incongruence effects – see below). While little is known about the role of the cerebellum in task-switching, this is certainly not the first study to find that it is significantly involved (De Bartolo *et al*., 2009; Dickson *et al*., 2016). It is perhaps not surprising that the brain structure most associated with motor control would be active during the resolution of motor conflict in a paradigm like the current one (Peterburs & Desmond, 2016). That it was not activated during isolated rule-switches (above) suggests that previous observations of cerebellar involvement in task-switch may have been confounding what might be thought of as “conceptual” rule-switches with motoric switches, an issue certainly worth pursuing more deeply in future studies.

We had predicted that the anterior cingulate cortex (ACC) would likely be involved in resolving the conflict created by incongruent stimuli, based on prior literature (Botvinick *et al*., 2001; Mayr, 2002; Kim *et al*., 2010). Here, however, it was for response-switching that we observed its involvement. It is important to note that the identified region here, while primarily situated in anterior cingulate, also extended somewhat anteriorly into medial frontal gyrus. Studies have consistently found the ACC involved in resolving incongruency of visual stimuli (Bush *et al*., 2000; Swainson *et al*., 2003; Fassbender *et al*., 2004; Woodward *et al*., 2006). One study also found anterior ACC involvement when there was incongruency in response mapping (i.e. response conflict) (van Veen & Carter, 2005), but it is difficult to interpret how this relates to our finding of deactivation of the ACC for a switch of response.

The postcentral gyrus also showed relative deactivation for response-switching compared to baseline, but it was more deactivated compared to response-repeating. And it is worth noting that although the identified region was primarily in postcentral gyrus, it did also extend anteriorly into precentral gyrus and posteriorly into inferior parietal cortex. The postcentral gyrus is known to receive and process somatosensory input (Iwamura, 1998; Iwamura *et al*., 2001; Staudt, 2010), but this cannot explain these findings because participants had their hand on the response pad for the duration of the study, and pressed the button equally in response-switch and response-repeat trials. However, another task-switching study, designed to limit working memory demands and isolate only switching processes, found postcentral gyrus involvement in task-switching, with the caveat that every task-switch also required a motor switch and thus this activation could be related to motor switch demands (Smith *et al*., 2004). Regarding the functional role that the postcentral gyrus may play during these switches, a study on motor reach errors found the postcentral gyrus to be involved in errors resulting in behavioral goal changes and not errors that lead to adaptation of limb dynamics (Diedrichsen *et al*., 2005). These findings suggest that the postcentral region is an area that performs and learns new visuo-motor transformations, which is relevant during switching of responses. However, it is important to understand why the current study found deactivation during response-switches instead of activation, as the above studies would suggest. While deactivation in functional neuroimaging studies is poorly understood overall, recently more studies have focused on deactivation, with a few specifically on motor deactivation (Allison *et al*., 2000; Kudo *et al*., 2004). Deactivation in motor cortices in response to motor activity has been shown, on the supposition that interhemispheric control of motor action also involves deactivation of the opposite hemisphere to prevent interference (Marchand *et al*., 2007). This fits well with our findings of response-switch deactivation only in the right hemisphere. All participants in this study used their right hand to respond to the task. Therefore, there could be right-hemisphere deactivation to allow for better left-hemisphere control, especially during the response-switch condition which requires altering the motor response and could therefore require increased processing.

The inferior temporal gyrus is part of the visual system’s ‘what’ pathway, such that lesions to this region result in impairments in object recognition (Creem & Proffitt, 2001). Response-switching may require more activation of the inferior temporal gyrus because there is increased need to interpret and categorize the stimulus before initiating a change in behavior. Cognitive flexibility has traditionally been thought of as a cortical process, occurring through frontal and parietal regions; however evidence indicates that subcortical regions are involved and even necessary (van Schouwenburg *et al*., 2010). The basal ganglia are thought to encode novel actions (Redgrave *et al*., 2013) and to help regulate response selection during cognitive flexibility tasks (Verstynen *et al*., 2012). Although there is little to go on from previous neuroimaging studies regarding response-switching, it would certainly be reasonable to expect activation in the striatum. In the context of task-switching, the striatum has been demonstrated to be active regardless of how the switch was cued (Liu *et al*., 2015). Patients with traumatic brain injury show a negative correlation between caudate body volume and task-switch accuracy, and a negative association between caudate head size and task-switch response times (Leunissen *et al*., 2014). As further evidence of the role of the basal ganglia, stroke patients with lesions focalized to the striatum had impaired task-switching performance (Cools *et al*., 2006). Alongside the striatum, the thalamus has also been shown to be engaged during cognitive flexibility (Liu *et al*., 2015). In a meta-analysis of cognitive control studies, the thalamus was significantly associated with inferior frontal junction (IFJ) activations, which is believed to be involved in the updating of the representation of the task. Since there is relatively little work on response-switching, a simple model of the respective roles of the identified regions might be useful. Based on our current findings and on the extensive related literature, the following processing sequence represents one plausible scenario: response-switching engages the inferior temporal gyrus for further visual processing of the stimulus, the middle frontal gyrus aids in decision-making based on the stimuli (Hare *et al*., 2009), the cerebellum detects a need to modulate the response (De Bartolo *et al*., 2009; Dickson *et al*., 2016), the striatum allows the conscious awareness of the new response requirement (Graybiel, 1998; Squire & Dede, 2015), and the thalamus relays that information back to the cortex to elicit the correct response (Saalmann *et al*., 2012; Bell & Shine, 2016).

### Stimulus incongruence

Examination of the effects of stimulus congruence (i.e. when the letter and number indicated the same motor response versus when they indicated conflicting motor responses) also revealed a network of regions that was more active during incongruent trials compared to congruent trials. Unique activations of the superior frontal gyrus, medial frontal gyrus, posterior cingulate, insula, and cerebellar regions were observed. Stimulus incongruence also activated a subset of regions that were recruited during the resolution of response-switching: middle frontal gyrus, inferior temporal gyrus, putamen, and two cerebellar loci. As mentioned above, posterior lobe cerebellar involvement was observed during the resolution of response-switches, as it was here during the resolution of stimulus-incongruence, but it was not observed when rule-switches were isolated. Again, it is perhaps unsurprising that the cerebellum might be specifically involved in situations where the conflict in the system is due to competing motor responses. In the case of response switches, this is likely because the currently indicated response does not match the previously executed one, and in the case of stimulus congruence, this may be because the currently indicated response does not match the contemporaneously cued response in the to-be-ignored stimulus dimension (e.g. what the letter is telling the participant to do while she or he is actively engaged in the number task).

Medial frontal gyrus is typically implicated in sustaining task-set representations (Cummings, 1995; Stuss *et al*., 2005), so it is perhaps surprising that this study revealed its involvement in resolving stimulus incongruence. This story is further complicated by the observation that one locus within the medial frontal gyrus showed more deactivation for incongruent stimuli compared to congruent stimuli, while another locus within medial frontal cortex showed increased activation for incongruent stimuli. The common observation here is that trials with incongruent stimuli elicited greater differential activity relative to baseline than did trials containing congruent stimuli; but perhaps the medial frontal gyrus has two distinct roles in resolving this conflict. It remains a puzzle, however, that we did not observe rule-switching related activation in this region, given its presumed role in task-set representation.

Insular cortex has been variously implicated in emotion regulation and the limbic system (Wager *et al*., 2004) as well as in multisensory processing (Bushara *et al*., 2003; Chen *et al*., 2015), but also during many studies that have investigated cognitive control processes (Fassbender *et al*., 2006; Simoes-Franklin *et al*., 2010; Droutman *et al*., 2015). Here, we observed insular involvement during processing of incongruent stimuli. One of the inherent qualities of incongruent stimuli is that one of the stimulus dimensions indicates a “go” response, whereas the other indicates a “stop” response (i.e. withhold or arrest your response). It is interesting therefore that in a previous study from our group, we observed clear insular activation for both cued and uncued STOPs in a variant of a response-inhibition task (Fassbender *et al*., 2009), and the current findings are also entirely consistent with a recent “activation likelihood estimation (ALE)” study that evaluated 111 neuroimaging studies that assessed neural activations in conflict-related paradigms (Li *et al*., 2017). These authors showed clear activation in bilateral insula in response to “stimulus-stimulus” conflicts; that is, tasks where the response demanded by the task-relevant feature of a stimulus was incompatible with the response associated with the task-irrelevant feature of the same stimulus, very much as was the case in the current study. Thus, it seems reasonable to propose that the insular activations we observe here are related to the need for motor suppression to the incompatible “instruction” provided by the irrelevant stimulus dimension (see Wessel and Aron, 2017 for a similar interpretation (Wessel & Aron, 2017)).

The posterior cingulate cortex was significantly involved in resolving trials with incongruent stimuli compared to congruent stimuli. The reader will recall that the PCC was also involved in rule-switches, but it is important to point out that the PCC region found active here for incongruent stimuli does not overlap with the regions identified during rule-switching. Studies have demonstrated distinct functional roles for different loci within the PCC, and the current findings clearly further support this multi-role interpretation (Leech *et al*., 2012; Liang *et al*., 2016). As with rule-switches, increased PCC activation during the resolution of stimulus-stimulus incongruence does not fit well with the notion that PCC should be less active during higher-load task performance (Gusnard *et al*., 2001; Raichle *et al*., 2001). Indeed, a recent study demonstrated greater PCC deactivation during a condition with incongruent stimulus-response mappings compared to congruent mappings (Li *et al*., 2015). This finding was interpreted as the need for suppression of task-irrelevant activity to allow for better external cognitive function, as might be expected during more taxing conditions (see (Anticevic *et al*., 2012)), and work has also shown that the strength of anti-correlation between the so-called default mode network (comprising PCC) and the fronto-parietal attentional control network is linked to better task performance in the form of less variable reaction times (Kelly *et al*., 2008). However, as with the current study, other studies have demonstrated a more “active” role for the posterior cingulate during taxing cognitive control operations, as we discussed above. What then is the role of the PCC in resolving the stimulus-stimulus incongruence inherent in the current task? One plausible explanation might lie in our use of orthographic stimuli. For example, in one recent study employing sentence stimuli, sentences were constructed such that they could either end with a word that was contextually congruent or one that wasn’t (e.g. “The pen leaked ink,” or “The pen leaked chocolate”). Increased PCC activation was observed for incongruent relative to congruent endings (Catarino *et al*., 2011), and other studies have also implicated the PCC in semantic processing (Frishkoff *et al*., 2004; Binder *et al*., 2009). Although our stimuli were considerably more basic letter-number combinations than the stimulus materials used in these studies, it is possible that the semantic decoding of them may have driven the increased PCC activation in the case of incongruent pairings.

We found the left thalamus involved in both rule-switching and incongruency, while the right thalamus was involved in response-switching. The thalamus has previously been implicated in task-switching (Liu *et al*., 2015), although its precise role is not clear. The thalamus is traditionally understood as a processing hub for sensory information (Nakajima & Halassa, 2017) and connecting subcortical regions to the cortex (Bell & Shine, 2016). It may also regulate cortical connectivity between networks for task completion (Nakashima *et al*., 2018), which fits well with the current study’s finding that there was thalamic involvement in all three sources of cognitive conflict. The thalamus is likely an important source for network control aiding changes in task sets or response sets.

The middle frontal gyrus, on the other hand, has been shown to be involved in reorienting attention and to be generally involved in attentional networks (Japee *et al*., 2015). Its involvement in resolving stimulus incongruence is consistent with the additional attentional demands that are likely required to discern the relevant stimulus dimension. Stimulus incongruence (and response-switching, as discussed above) also activated the putamen in the current study. The putamen has been shown to aid in rule-based learning (Ell *et al*., 2006), and so this finding further supports its role in category learning and decision-making tasks. Another necessary process during task-switching studies is to appropriately process and interpret the visual stimuli. The inferior temporal gyrus is integral to this process (Denys *et al*., 2004) and was clearly observed in the current study for trials with incongruent stimuli, which can be reasonably expected to require increased visual processing compared to trials with congruent stimuli. Many studies have demonstrated that the ACC plays a role in the error monitoring and detection necessary to reallocate attention (Botvinick *et al*., 2001; Mayr *et al*., 2003; Swainson *et al*., 2003; Fassbender *et al*., 2004; Derrfuss *et al*., 2005; Liston *et al*., 2006; O’Connell *et al*., 2007; Kim *et al*., 2010). Although the current study’s cingulate activation was not as anterior as seen in many of these prior studies, it likely reflects a similar functional role. Studies that have tried to determine whether ACC is involved in stimulus-only conflict or response conflict have found that ACC is involved when there is a conflict in the response (Milham *et al*., 2001; Liu *et al*., 2006). This is in line with our results because the irrelevant stimulus in a given trial was incongruent because it elicited a different behavioral response than the relevant stimulus. ACC activation can be modulated by top-down motivation or awareness. Greater ACC activation has been linked to awareness of errors, compared to unaware errors (Orr & Hester, 2012), and motivation through incentives can differentially activate rostral ACC compared to caudal ACC (Simoes-Franklin *et al*., 2010). The current findings of ACC activation further cement its role as a monitor of potential conflict and need for regulation in task-switching studies, but likely in other domains as well. The superior frontal gyrus is often found as an integral region during task-switching (Nagahama *et al*., 1999; Cutini *et al*., 2008). Because of its role in resolving stimulus incongruence in the current study, we surmise that it may have a more general role in resolving cognitive conflict that allows for the filtering of irrelevant stimuli and engagement of the appropriate task.

### Study Limitations

A limitation of this study is the relatively modest number of participants in our cohort, raising issues of power and generalizability. Although 11 participants were studied, it merits re-emphasizing that this study comprised a total of 42 scan sessions, with the express purpose of this design being to take advantage of a repeated measures statistical approach in analyzing both the neuroimaging and behavioral data. It also bears emphasizing that within each of the 42 scanning sessions, the density of data was considerably greater than is typical of the great majority of studies in this field, where task-related data for each participant are typically derived from sessions on the order of 10-20 minutes each. Here, each participant provided upwards of 3.5-to-4 hours of data for the task-based analysis, more than an order of magnitude more data than is typical. When considering how well a study is powered, the density of data available for each subject is a key consideration. In all experimental designs, the goal is to optimize the investigator’s ability to observe possible signal relative to background noise. While increasing sample size is one way to do this, another equally effective way is to increase the number of repeated measures made of any given item/participant, as was done here. Generalizability is a concern with any study that does not adequately sample a representative group of individuals and this is certainly also the case with this study where the participants represent a convenience sample, specifically recruited because they were easily accessible and could be relied upon to return for repeated experimental sessions.

Another potential limitation pertains to the relatively rapid and regular stimulus presentation rates utilized here and how these might affect the ability of the applied linear model to effectively extract processes that were exclusively related to the condition of interest without other collinearities. Of course, in testing task-switching processes, it is key that relatively fast and consistent cue-to-target intervals are used so that the processes of interest are appropriately taxed. This is because the use of larger inter-trial-intervals introduces entirely new factors, such as differential time-based task-set decay, while varying the length of inter-trail-intervals has been shown to introduce sequential dependencies, such that switch costs on a current trial are affected not only by the inter-trial-interval leading into that trial, but also by the one preceding it (Grange & Cross, 2015). Therefore, the temporal structure of the current paradigm was designed to eliminate these other confounds and was a necessary compromise. Returning to the original concern regarding the potential for collinearities, it is also key to point out that while the alternation between trials was rapid and consistent in the current study, the occurrence of the each of the conditions of interest was in fact random. As such, the timing between instances of a given condition of interest (e.g. rule-switches) varied from 2 seconds to 60 seconds (see Supplementary Figure E), a temporal structure that lends itself very well to the event-related fMRI approach.

As with any study, there are some limitations of interpretation based on our study design. One important aspect to note is that every time there was a rule-switch, there was also a cue-switch (i.e. going from a blue rectangle to a yellow rectangle). This can be a possible confound in that we have not isolated rule-switch effects *independent* of cue-switch effects. Part of this issue is addressed because rule-switch trials included both those that had blue cues preceded by yellow cues and trials that had yellow cues preceded by blue cues. However, one study that did explicitly investigate cue-switch processes alone found that lateral premotor cortex, inferior temporal gyrus, and fusiform gyrus were involved in encoding the cue information (Brass & von Cramon, 2004). We did not find these regions in our rule-switch analysis and therefore feel that it is unlikely that we are capturing activity purely due to cue-switch effects.

Lastly, in the conjunction analyses presented herein, some of the overlapping voxels are limited to a relatively small area. Unfortunately, this study does not allow us to determine whether such instances might represent two distinct but overlapping loci of activity within the same region or whether these represent simple activations of the same region. Here, we have chosen to discuss these findings as brain regional activations, and it will fall to future work to more precisely investigate specific overlap patterns within a given functional region.

### Conclusion

This study demonstrates substantial differences in network activation for three distinct aspects of cognitive flexibility within a single task-switching paradigm, pointing to largely dissociable sources of competition (i.e., mostly non-overlapping neural circuits) as the primary modus by which separable sources of competition are resolved. We did not find evidence for any one set of regions that was common to all three sources of competition, although some regions were found to participate in conflict resolution across two of the sources. Thus, while the study does not point to a set of domain-general processes, it does suggest shared circuitry for some aspects of conflict resolution, with regions such as the precuneus, middle frontal cortex, and inferior temporal gyrus implicated in resolving more than one source of conflict during task performance.

## Acknowledgments

The authors thank Ms. Sarah Fendrich of Ossining High School who provided valuable assistance during a summer internship with KMB at the Einstein Cognitive Neurophysiology Laboratory.

## Data Sharing Statement

At the time of publication, the authors will make all relevant data from this project available on a publicly accessible data repository (e.g. Figshare).

## Research involving Human Participants and/or Animals

This research involved the participation of human subjects. The Institutional Review Board of Albert Einstein College of Medicine approved all materials and procedures, and all ethical guidelines were in accordance with the tenets of the Declaration of Helsinki.

## Informed Consent

Informed, written consent was obtained from all participants included in this study.

---

**Supplementary Table A.** Acronyms Used in This Paper.

|  |  |
| --- | --- |
| ACC | anterior cingulate cortex |
| ASD | autism spectrum disorders |
| DLPFC | dorsolateral prefrontal cortex |
| EEG | electroencephalography |
| fdr | false discovery rate |
| fMRI | functional magnetic resonance imaging |
| IFJ | inferior frontal junction |
| PPC | posterior parietal cortex |
| pre-SMA | pre-supplementary motor area |
| RFX-GLM | random-effects general linear model |
| RT | reaction time |
| TE | echo time |
| TR | repetition time |

**Supplementary Figure A.**
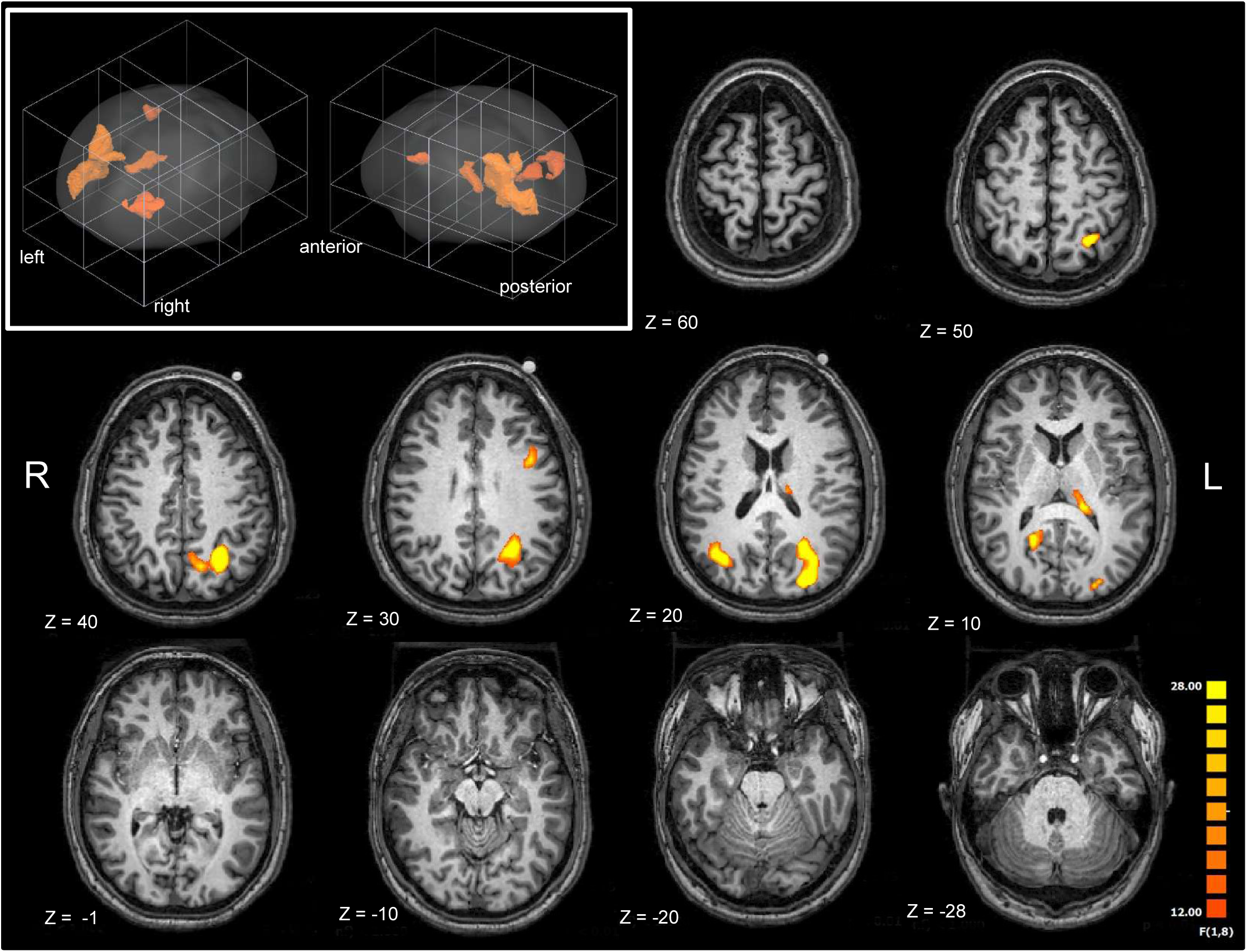
Map of regions significantly involved in rule-switching excluding the one potential outlier participant. The upper left panel shows all the significant regions in orange in a 3D rendering. In the horizontal slices, the color range is according to the F-stat for each voxel, ranging from 12 to 28. Images are in radiological format (right is left), and z coordinates are in Talairach space. This map shows a strong resemblance with Figure 3 with all participants.

**Supplementary Figure B.**
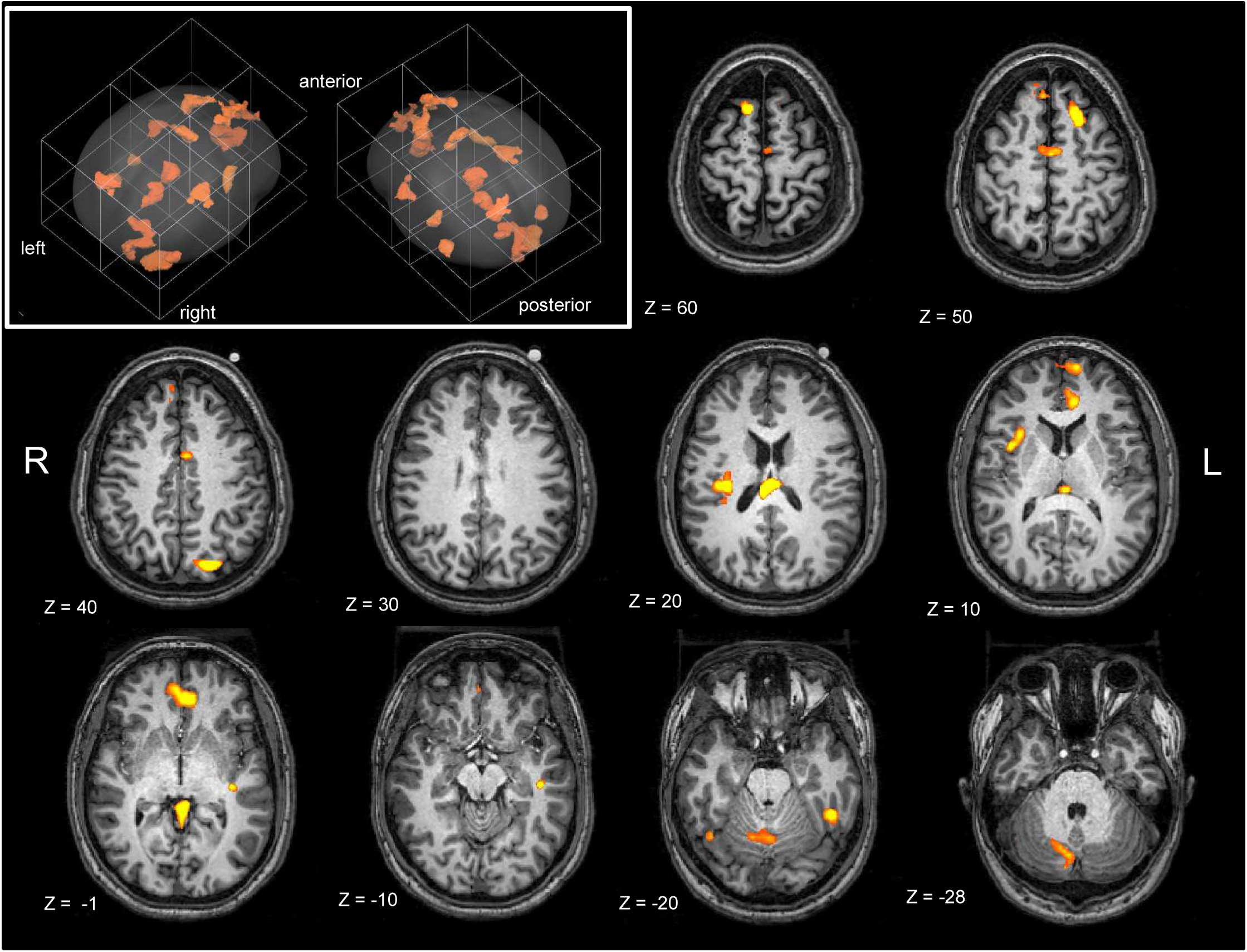
Map of regions significantly involved in incongruency, excluding one potential outlier participant. The upper left panel shows all the significant regions in orange in a 3D rendering. In the horizontal slices, the color range is according to the F-stat for each voxel, ranging from 12 to 28. Images are in radiological format (right is left), and z coordinates are in Talairach space. This map is very similar to Figure 4 which contained all participants together.

**Supplementary Figure C.**
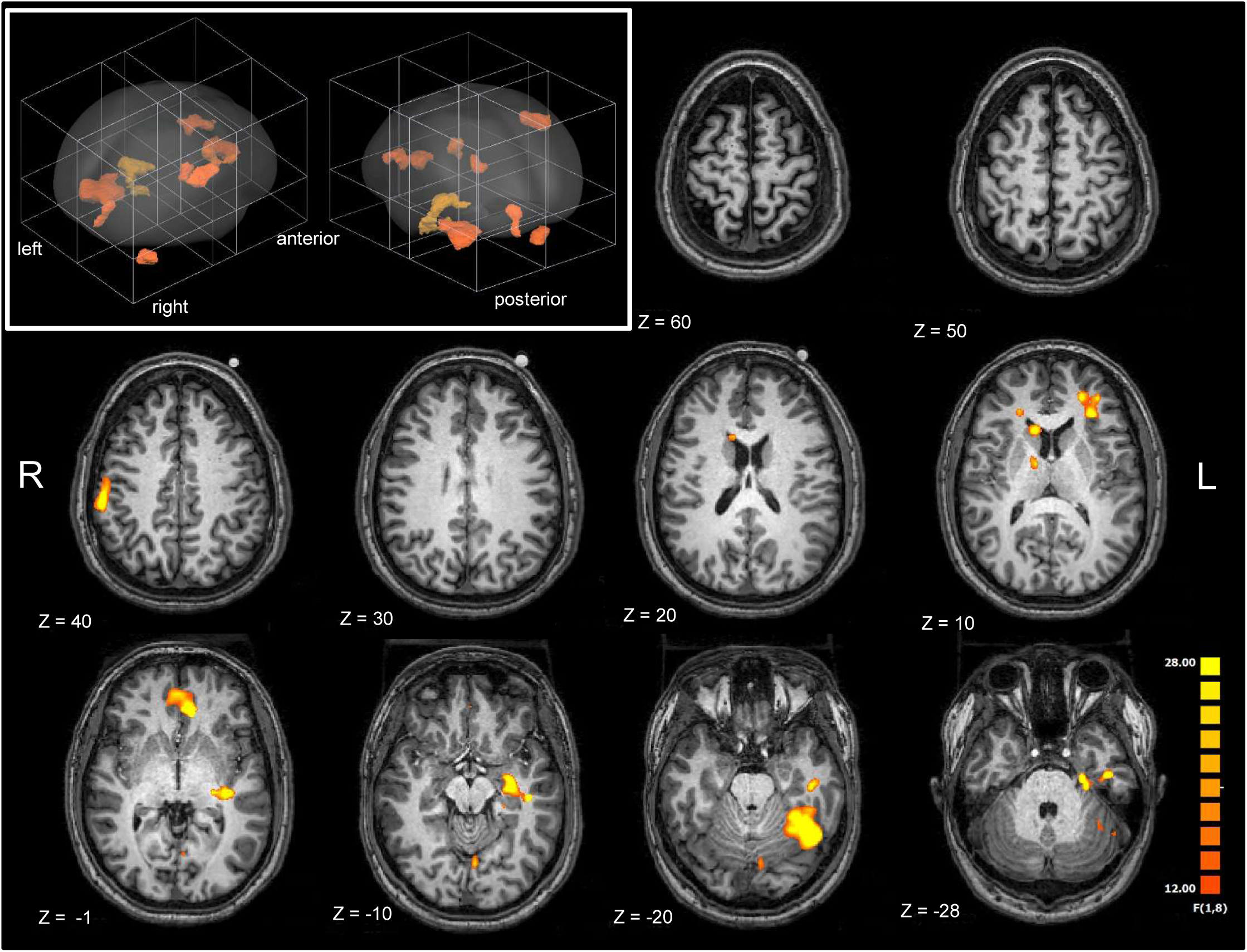
Map of regions significantly involved in response-switching, excluding one potential outlier participant. The upper left panel shows all the significant regions in orange in a 3D rendering. In the horizontal slices, the color range is according to the F-stat for each voxel, ranging from 12 to 28. Images are in radiological format (right is left), and z coordinates are in Talairach space.

**Supplementary Figure D.**
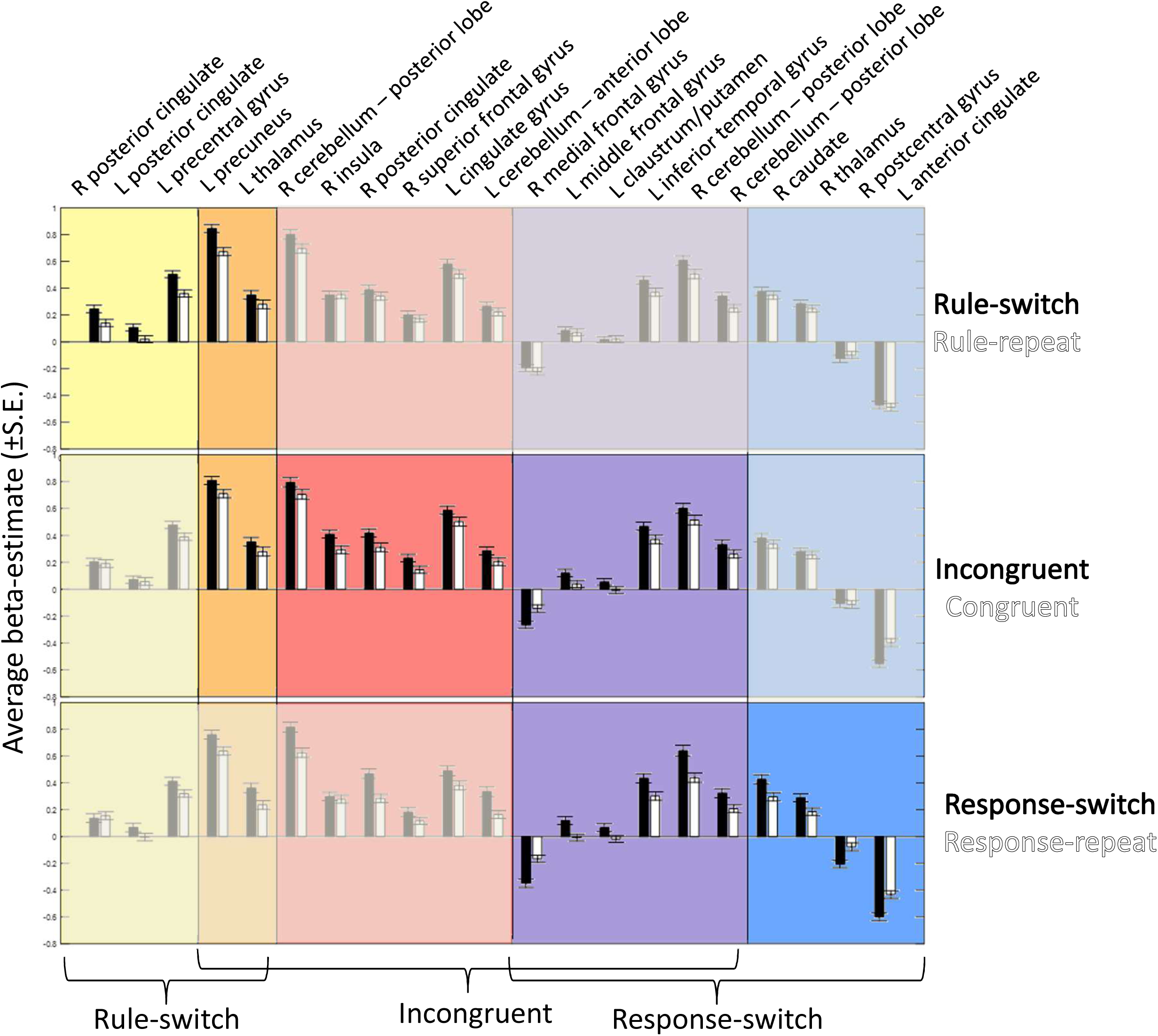
Beta-estimates for each significant cluster found in our analyses under each condition. They are sorted by which analyses they came out significant for. The top row shows beta-estimates for rule-switching and rule-repeating, the middle row is for incongruent and congruent trials, and the bottom row shows estimates for response-switching and response-repeating. Averages and standard errors are computed with the beta-estimates from each run from each participant.

**Supplementary Figure E.**
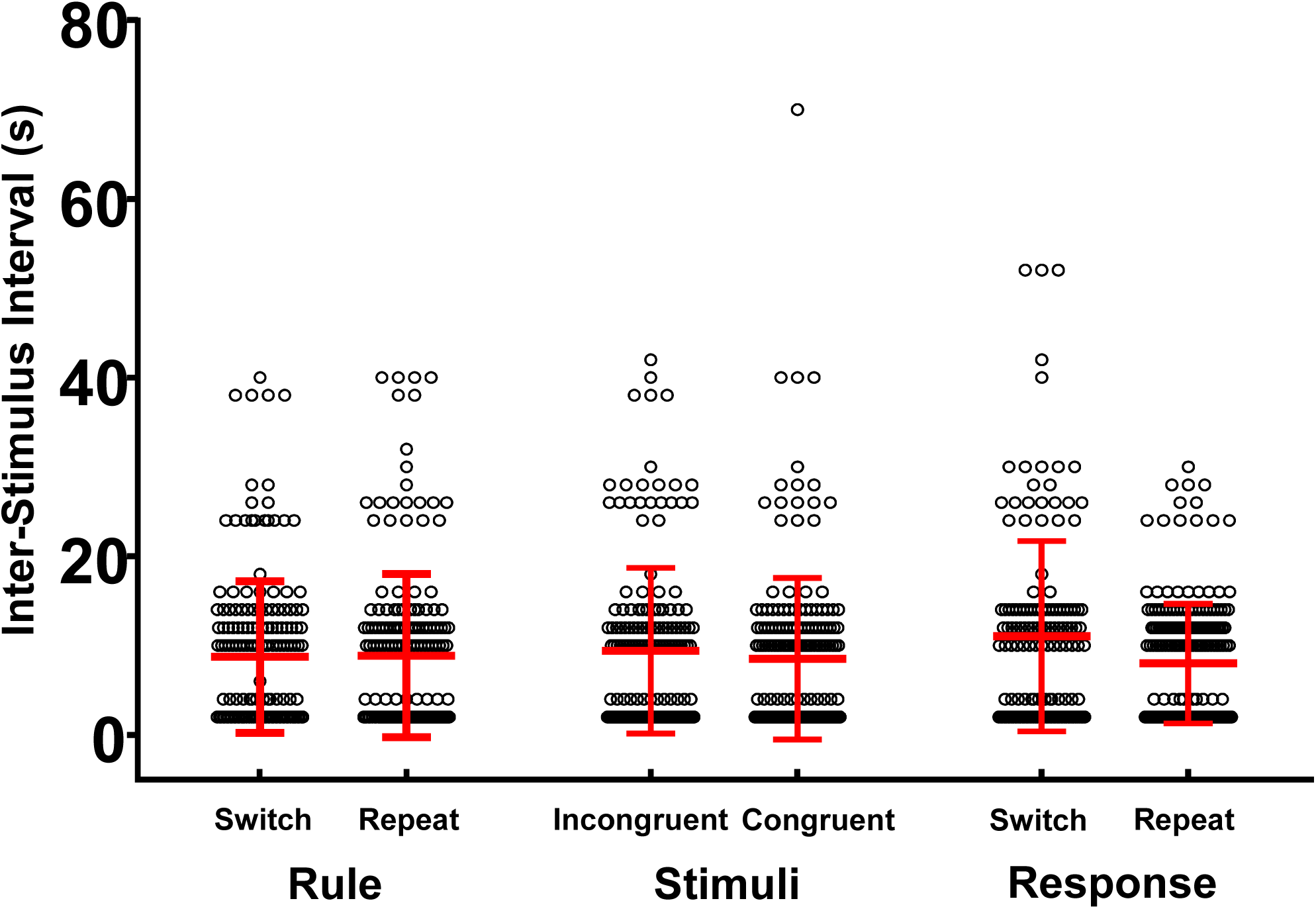
Event-related fMRI studies can succumb to an issue where the conditions occur at the same time throughout runs or scan sessions and confound the general linear model. Here we demonstrate that the conditions in our study were randomly placed through each run and session so that, although the average inter-stimulus interval is 9 seconds, the standard deviation and variability is similarly around 10 ms.

